# A Bayesian analysis of the association between *Leukotriene A4 Hydrolase* genotype and survival probability of tuberculous meningitis patients treated with adjunctive dexamethasone

**DOI:** 10.1101/2020.08.03.233791

**Authors:** Laura Whitworth, Jacob Coxon, Arjan van Laarhoven, Nguyen Thuy Thuong Thuong, Sofiati Dian, Bachti Alisjahbana, Ahmad Rizal Ganiem, Reinout van Crevel, Guy Thwaites, Mark Troll, Paul Edelstein, Roger Sewell, Lalita Ramakrishnan

## Abstract

Tuberculous meningitis (TBM) remains the most devastating form of tuberculosis (TB) with high mortality despite effective antimicrobial treatment. As mortality has been linked to excessive inflammation, anti-inflammatory glucocorticoids are now routinely used as adjunctive treatment with antimicrobial therapy. However, they reduce mortality by only ~ 30%, raising the possibility that only a subset of TBM deaths are caused by inflammatory pathophysiology. Studies in Vietnam found that the survival benefit of adjunctive glucocorticoids was limited to individuals with a common promoter variant in the *leukotriene A4 hydrolase* (*LTA4H*) gene encoding an enzyme that regulates inflammatory eicosanoid expression. The variant constitutes a C/T transition with TT homozygotes having increased expression over CT heterozygotes and CC homozygotes. In Vietnam, the *LTA4H* TT genotype predicted survival, consistent with dexamethasone benefiting only those individuals with a dysregulated hyper-inflammatory response. However, a study of TBM patients in Indonesia did not find the *LTA4H* TT genotype to confer a significant survival benefit. Given the potential of personalized life-saving anti-inflammatory therapies guided by *LTA4H* genotype, we have used Bayesian methods to analyze the data from both studies. Bayesian analysis reveals that the *LTA4H* TT genotype confers survival benefit in both the Vietnam and Indonesia cohorts that begins within days and continues long-term. However, its benefit is nullified in the most severe cases where other factors cause early mortality. *LTA4H* TT genotype is associated with increased survival in HIV-positive patients also. Thus, our analysis extends the association of *LTA4H* genotype with TBM survival to populations outside of Vietnam and to HIV-positive patients. Patient *LTA4H* genotyping used in conjunction with disease severity assessment may help to target glucocorticoids to patients most likely to benefit from this broadly-acting immunosuppressive regimen despite its significant adverse effects.

## INTRODUCTION

Tuberculous meningitis (TBM) is the most severe form of tuberculosis. Despite effective antimicrobial therapy, it results in 20-25% mortality in HIV-negative individuals and ~ 40% mortality in HIV-positive individuals (Stadelman et al., 2020, in press; Thwaites, van Toorn, & Schoeman, 2013). A long-standing hypothesis that an excessive intracerebral inflammatory response underlies TBM mortality (Shane & Riley, 1953) led to multiple trials of adjunctive anti-inflammatory treatment with glucocorticoids (e.g. dexamethasone) (Prasad, Singh, & Ryan, 2016; Wilkinson et al., 2017). Findings from a randomized controlled trial (RCT) in Vietnam that adjunctive dexamethasone improved survival by ~ 30% led to it becoming standard of care treatment (Thwaites et al., 2004). However, the modest benefit of adjunctive dexamethasone treatment suggested a heterogeneity in glucocorticoid-responsiveness (Donald & Schoeman, 2004; Schoeman & Donald, 2013). Findings in a zebrafish model of TB provided a testable hypothesis for a mechanism underlying this heterogeneity (Thwaites et al., 2004; Tobin et al., 2012; Tobin et al., 2010). The zebrafish findings suggested that either deficiency or excess of leukotriene A4 hydrolase (LTA4H), a key enzyme that regulates the balance of pro- and anti-inflammatory eicosanoids, increase susceptibility to TBM for opposite reasons - too little or too much inflammation (Tobin et al., 2012; Tobin et al., 2010). It became possible to test the prediction when a common human functional *LTA4H* promoter variant (rs17525495) was identified comprising a C/T transition that controlled LTA4H expression, with the T allele causing increased expression (Tobin et al., 2012). A retrospective analysis of patient *LTA4H* rs17525495 genotypes in the Vietnam dexamethasone RCT cohort confirmed the prediction (Thwaites et al., 2004; Tobin et al., 2012). Among HIV-negative patients, the survival benefit of dexamethasone was restricted to patients with the hyper-inflammatory TT genotype, with CC patients potentially harmed by this treatment (Tobin et al., 2012). These findings supported the model that mortality from TBM was due to two distinct inflammatory states, and that *LTA4H* genotype might be a critical determinant of inflammation and consequently of the response to adjunctive anti-inflammatory treatment. If true, then personalized genotype-directed adjunctive glucocorticoid treatment would be warranted, with the drug given only to TT patients. This would be particularly important given the possible harm to the hypo-inflammatory CC group, as well as the adverse effects of long-term high dose treatment with a broadly acting immunosuppressant.

To further these findings, two new studies of the association of *LTA4H* genotype with TBM survival in HIV-negative patients were performed in Vietnam and Indonesia, respectively (Thuong et al., 2017; van Laarhoven et al., 2017). Because glucocorticoid adjunctive therapy had become standard of care owing to the benefit observed in the randomised controlled trial (Thwaites et al., 2004), all patients received it in both studies. Therefore, the prediction that could be tested was that TT mortality is less than CC+CT mortality. Whereas the Vietnam study confirmed this prediction, the Indonesia study did not. The Vietnam cohort had an overall mortality of 18.8%, similar to that reported in the literature (Stadelman et al., 2020, in press). A striking feature of the Indonesia cohort was its more than two-fold increased mortality in comparison with the Vietnam cohort. Moreover, most of the Indonesia cohort deaths occurred early with a median time to death of eight days versus 50 days in Vietnam (Table 1). This high early mortality raised the possibility that the impact of the *LTA4H* variant differs by disease severity, and may not be relevant in more severe disease (Fava & Schurr, 2017). If so, then the effects of the *LTA4H* genotype were being masked by the preponderance of extremely severe cases in the Indonesia cohort (Fava & Schurr, 2017).

**Table 1.** Characteristics of the Vietnam and Indonesia cohorts.

|  | **HIV-negative** |  | **HIV-positive** |
| --- | --- | --- | --- |
|  | **Vietnam** | **Indonesia** | **Vietnam** |
| **Total** | 439 | 376 | 325 |
| **Glasgow Coma Score**<br>**median (range)** | 15 (4-15) | 13 (5-15) | 15 (3-15) |
| **BMRC TBM grade**<br>**no. (% of total)** |  |  |  |
| **1** | 163 (37.1) | 34 (9.0) | 137 (42.15) |
| **2** | 206 (47.0) | 284 (75.5) | 124 (38.15) |
| **3** | 70 (15.9) | 58 (15.4) | 64 (19.7) |
| **Age in years**<br>**median (range)** | 41 (18-93) | 28 (14-90) | 33 (18-63) |
| **Age in years by TBM grade**<br>**median (range)** |  |  |  |
| **1** | 39 (18-85) | 27 (16-45) | 32 (18-53) |
| **2** | 47 (18-93) | 29 (14-90) | 34 (20-63) |
| **3** | 33 (18-86) | 26 (14-64) | 33 (19-46) |
| **Overall mortality**<br>**no. (%)** | 83 (18.8) | 146 (39.9) | 129 (40.4) |
| **Time to median mortality**<br>**(days)** | 50 | 8 | 37 |
| **Mortality by BMRC TBM grade**<br>**No. dead (% of grade)** |  |  |  |
| **1** | 12 (6.8) | 5 (15.9) | 32 (23.8) |
| **2** | 45 (21.9) | 106 (38.0) | 50 (40.9) |
| **3** | 26 (37.9) | 35 (63.8) | 47 (75.2) |

Both studies used, as the primary metric of significance testing, Cox regression modelling, an approach that assumes that the ratio of hazard rates between groups is constant throughout the observed period (Bradburn, Clark, Love, & Altman, 2003; Greenland et al., 2016). Therefore, it could miss important differences in these studies of TBM, a disease which can present acutely yet have a prolonged time course with vastly differing mortality risks over time (Thuong et al., 2017; Thwaites et al., 2013; van Laarhoven et al., 2017). Moreover, testing the hypothesis that *LTA4H* effects are limited to specific disease severity grades requires subgroup analysis. The use of frequentist statistics would limit the ability to perform such subgroup analyses because the penalties it sets for multiple comparisons do not reflect real-world situations (Gelman & Loken, 2013; Smith & Ebrahim, 2002). Bayesian analysis is ideally suited to simultaneously estimate treatment effects in multiple subgroups because it results in different interactions with the number of results obtained which are much less problematic than those arising in frequentist analysis (Box 1).

Therefore, we used a Bayesian approach to analyze data from the two cohorts (Gelman & Loken, 2013; MacKay, 2003; Zampieri et al., 2020) (see also Methods). Bayesian analysis also enables the detection of significant differences that might be limited to just a part of the time-course and therefore would allow analysis to be independent of the kinetics of death in the Vietnam and Indonesia cohorts. Finally, medical management decisions are guided by an assessment of the probabilities of outcome. In TBM, the question faced by the clinician is how likely is glucocorticoid therapy going to help or harm a patient. Bayesian paradigms, unlike frequentist ones, understand probability in a real-world way, using it to indicate the plausibility of a particular conclusion (MacKay, 2003; Zampieri et al., 2020).

The severity grade-specific analyses, coupled with temporal analyses made possible by Bayesian methods, reveal that the *LTA4H* TT genotype is associated with survival in both HIV-negative cohorts and that this association extends to HIV-positive patients as well.

## METHODS

The anonymized patient cohort data used here has been previously described in detail (Thuong et al., 2017; van Laarhoven et al., 2017). Our analysis included all 439 HIV-negative patients from the Vietnam study and 376 of the 515 HIV-negative patients from the Indonesia study. The remaining 139 Indonesian patients were excluded for lack of information on *LTA4H* rs17525495 genotype (87), baseline disease severity (44) or outcome (8). All 325 HIV-positive patients from Vietnam were analyzed. HIV-positive patients from Indonesia were excluded due to small sample size and lack of the information above (n=93). All Vietnam patients were admitted to one of two tertiary care referral hospitals in Ho Chi Minh City, Vietnam: Pham Ngoc Thach Hospital for Tuberculosis and Lung Disease (designated Hospital 1) and the Hospital for Tropical Diseases (designated Hospital 2) (Heemskerk et al., 2016). All patients were treated with adjunctive dexamethasone for the first 6-8 weeks with the regimen adjusted to disease severity on presentation (Thwaites et al., 2004).

Patient cohorts were compared overall as well as stratified into disease severity groups based on the TBM grade and by *LTA4H* genotypes, into the TT group (previously linked with response to steroids) and the combined CC and CT genotypes (non-TT group). The Bayesian analysis methods used are detailed in Supplementary Methods. We limited analysis to the first 9 months of the one-year observation period in Indonesia to be compatible with the 9-month observation period in Vietnam.

## RESULTS

### Characteristics of the Vietnam and Indonesia HIV-negative cohorts

The age range of patients was similar in the Vietnam and Indonesia cohorts with Indonesia patients tending to be younger (Table 1). We compared the cohorts for disease severity on presentation using both measures used in the studies, the Glasgow Coma Score (GCS) and the modified British Medical Research Council TBM grade (TBM grade) (Box 2). The Indonesia cohort had more severe disease on presentation by both measures (Table 1 and Figure S1). We used the TBM grade for further analyses because it divides patients into just three severity groups, making comparisons more feasible. Importantly, it also provides clinically relevant separation of GCS 15 patients, the most highly represented in both cohorts (Figure S1), into those with and without focal neurological signs.

### LTA4H TT genotype association with survival becomes stronger with increasing disease severity in Vietnam HIV-negative patients

Because the Indonesia cohort was skewed towards more severe disease on presentation, one explanation for the lack of an *LTA4H* genotype association with survival in Indonesia was that the underlying association is overridden by severe disease, a strong independent correlate of mortality (Fava & Schurr, 2017; Wang et al., 2019). Indeed, a detailed comparison of the Vietnam and Indonesia cohorts showed that 76% of Indonesia patients presented in Grade 2 versus 47% of the Vietnam patients (Table 1). This increase was driven entirely by a shift from Grade 1 (9% vs 37% in Vietnam). The cohorts had nearly equal proportions of Grade 3 patients (15% each). Therefore, the ~ 2-fold-increased overall mortality in Indonesia could be largely accounted for by a corresponding increase of Grade 2 patients (1.6 fold higher than Vietnam). Disease severity as assessed by BMRC Grade or GCS at presentation is also a strong predictor of earlier death (Fava & Schurr, 2017; Wang et al., 2019), and the Indonesia patients died sooner (median time to death 8 days versus 50 days in Vietnam) (Table 1).

If *LTA4H* genotype associates with survival most strongly in mild disease, then the association seen for the entire Vietnam cohort (Thuong et al., 2017) should be strongest in Grade 1 patients. We tested this prediction with Bayesian analysis using a prior that was intentionally uninformative and very wide, while still being centered on clinician-expected survival curves and hazard rates. We included additional parameters to allow for the possibility that not all patient deaths would be linked to the same mode of death. Importantly, the model and priors used allowed us to incorporate our pre-existing knowledge that mortality risk to a population of TBM patients varies smoothly with time, rather than occurring at a number of discrete times common to all patients as is implied by the maximum likelihood solution illustrated by a Kaplan-Meier plot. The details of the model and the priors are in Supplementary methods. The definitions of terms and abbreviations used throughout the paper are in Box 3.

In the original Vietnam study, the TT genotype was associated with survival and the CC and CT genotypes had similarly increased mortality over TT (Thuong et al., 2017), so we compared TT survival to that of CT and CC combined (non-TT). We first confirmed that the TT variant distribution did not differ by grade on presentation (Table 2). Bayesian analysis confirmed that the TT genotype was associated with a survival advantage in the overall cohort (Figure 1A). Moreover, the analysis revealed that this survival advantage manifested early and persisted over most of the observation period (Figure 1A). The Bayesian method enables a more detailed evaluation of mortality risk over time through a hazard rate analysis. This analysis reinforced the significantly higher hazard rate for non-TT starting at 4 days and persisting through 120 days (Figure 1A, inset).

**Figure 1.**
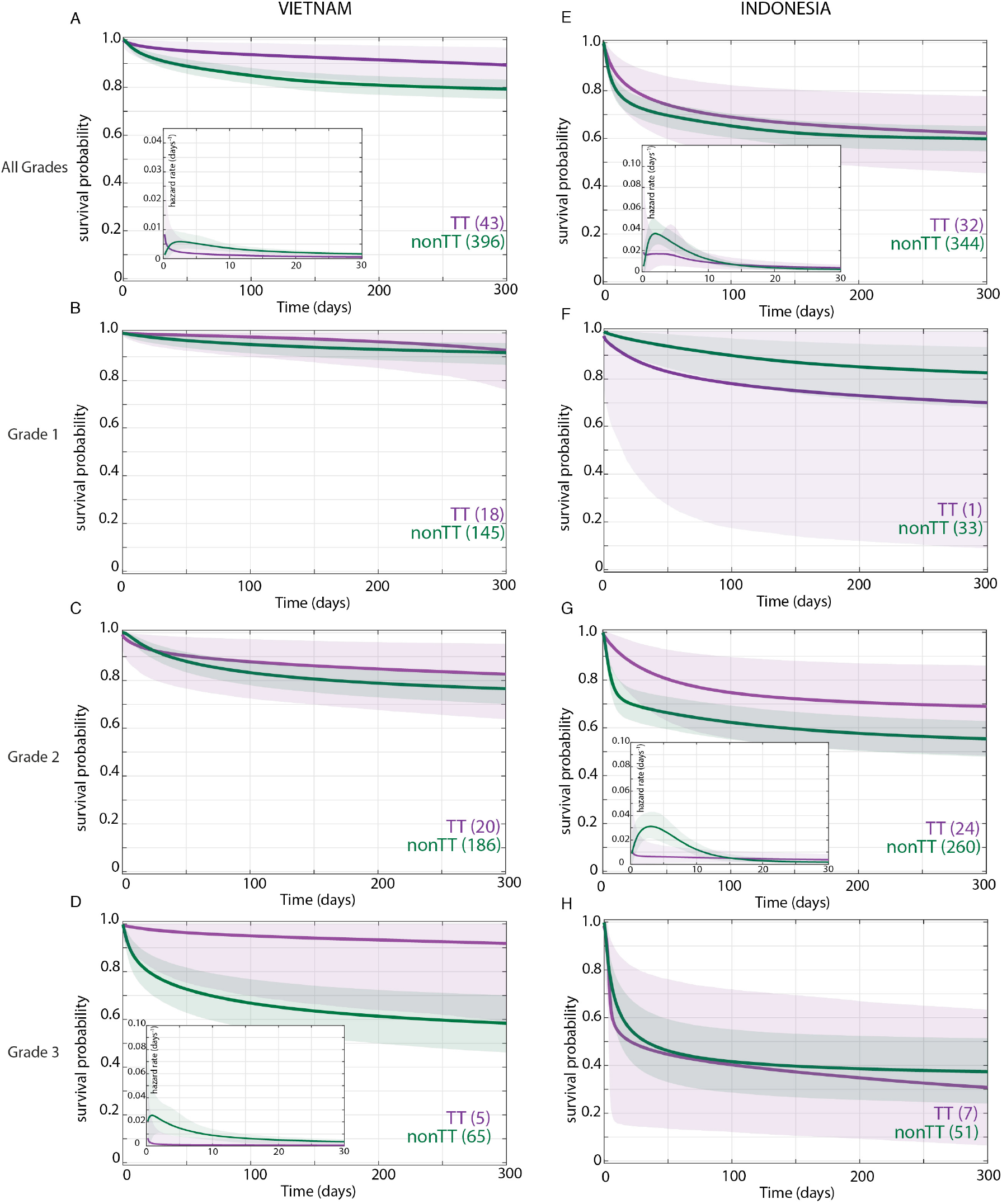
Effect of LTA4H rs17525495 genotype on HIV-negative patient survival probability over all grades in Vietnam (A) and Indonesia (E), and stratified by grade (B - D, F - H). Coloured lines represent mean posterior survival probability curves for the nine-month observation period. Shaded areas represent 95% Bayesian confidence limits for posterior probability. Comparisons where TT (purple) to non-TT (green) differences were significant have boxed insets showing hazard rates for the first 30 days; all other comparisons, not significant. The number of patients at the starting time point are indicated in parentheses. In Vietnam, overall (A), TT survival was significantly higher than non-TT from day 39 onwards with maximum probability 0.98, survival gap 11%; non-TT hazard rate was significantly higher than TT from day 4 to day 120, with their ratio peaking at 3 on day 6 and remaining >1 until day 223. (D) Grade 3 TT survival was significantly higher from day 3 onwards with maximum probability 0.97, survival gap 30%. The TT hazard rate dropped from the start, while the non-TT hazard peaked at 16 times higher than TT on day 3; non-TT over TT hazard rate ratio remained > 1 throughout. In Indonesia, overall (E), TT survival was non-significantly higher than non-TT (maximum probability 0.92); the non-TT hazard rate was greater than the TT hazard rate from day 1 to day 13, significantly so (and by 2-fold) on days 2 and 3 (maximum probability 0.97). (F) Grade 1 comparisons were uninformative due to TT sample size (n=1). (G) Grade 2 TT survival was significantly higher on days 4-32 with maximum probability 0.99, survival gap 9%. The TT hazard rate dropped from the start, while the non-TT hazard peaked at 5 times higher than TT on day 3. The non-TT over TT hazard rate ratio remained > 1 until day 15.

**Table 2.** LTA4H genotype frequency in HIV-negative and HIV-positive cohorts.

|  | **HIV-negative** |  | **HIV-positive** |
| --- | --- | --- | --- |
|  | **Vietnam** | **Indonesia** | **Vietnam** |
| **rs17525495 LTA4H**<br>*no. (% total)* |  |  |  |
| **CC** | 184 (41.9) | 216 (57.5) | 146 (44.9) |
| **CT** | 212 (48.3) | 128 (34.0) | 133 (40.9) |
| **TT** | 43 (9.8) | 32 (8.5) | 46 (14.2) |
| **No. (% of grade total)** |  |  |  |
| **Grade 1** |  |  |  |
| **CC** | 64/163 (39.3) | 21/34 (61.8) | 59/137 (43.1) |
| **CT** | 81/163 (49.7) | 12/34 (35.3) | 56/137 (40.9) |
| **TT** | 18/163 (11.0) | 1/34 (2.9) | 22/137 (16.1) |
| **Grade 2** |  |  |  |
| **CC** | 86/206 (41.8) | 161/284 (56.7) | 57/124 (46.0) |
| **CT** | 100/206 (48.5) | 99/284 (34.9) | 52/124 (41.9) |
| **TT** | 20/206 (9.7) | 24/284 (8.5) | 15/124 (12.1) |
| **Grade 3** |  |  |  |
| **CC** | 34/70 (48.6) | 34/58 (58.6) | 30/64 (46.9) |
| **CT** | 31/70 (44.3) | 17/58 (29.3) | 25/64 (39.1) |
| **TT** | 5/70 (7.1) | 7/58 (12.1) | 9/64 (14.1) |

When we stratified the Vietnam patients by grade and *LTA4H* genotype, we got a surprising result. The *LTA4H* TT association with survival was barely present in Grade 1, a bit more in Grade 2, and strongest in Grade 3 where it reached significance (Figure 1B-D). Similar to the overall cohort, the Grade 3 increased survival probability for TT also manifested early and persisted throughout (Figure 1D). Hazard rate analysis again showed that non-TT patients had a greatly increased risk of mortality very early (Figure 1D, inset). The non-TT over TT hazard rate ratio peaked at 16 on day 3 (Figure 1D, inset). This early high peak dropped only gradually over time; it was 5 at day 100 and remained >1 throughout (data not shown).

In sum, our analysis revealed that in Vietnam, *LTA4H* TT was associated with survival, not in mild disease as suggested earlier (Fava & Schurr, 2017), but rather in the most severe disease grade. In fact, the bulk of the overall association was being driven by Grade 3 patients who constituted only 15.9% of the cohort (Table 1). Non-TT patients were at greatest risk of dying within days of admission, a risk that diminished with time but remained greater than the TT patients throughout.

### In Indonesia HIV-negative patients, the LTA4H TT genotype effect does not extend beyond Grade 2

Bayesian analysis found that, in the overall Indonesia cohort, survival of the TT patient group was higher than non-TT though falling short of significance (maximum probability 0.92) (Figure 1E). Moreover, this analysis detected that the hazard rate for non-TT patients was higher than TT patients from day 1 to day 13; the non-TT over TT ratio reached significance on days 2 and 3, at which time the non-TT hazard rate was twice that of TT. Thus, while the TT beneficial effect was weaker than in Vietnam (compare Figure 1E to 1A), hazard rate analysis showed that as in Vietnam, TT benefit manifested early (Figure 1E inset, compare to Figure 1A inset). For the grade stratified cohorts, the analysis of Grade 1 patients was uninformative as there was only one TT patient in this group who survived throughout (Table 2 and Figure 1F; also see Supplementary Methods section 4). In Grade 2, the TT survival effect was significant, in contrast to Grade 2 Vietnam (compare Figure 1G to 1H). Rather, the pattern of the Grade 2 association was similar to Vietnam Grade 3 with a significant early TT survival benefit. As in Vietnam, the TT survival benefit started within days with an early hazard rate peak for the non-TT group. In Grade 3, the *LTA4H* TT effect was again absent.

Since Grade 2 patients constitute the bulk of the Indonesia cohort (75.5%), why was the *LTA4H* genotype effect in this grade not reflected in the overall cohort analysis? This was particularly curious given that in Vietnam the overall significant effect was being driven very substantially by Grade 3 patients who constituted only 15.9% of the cohort. We saw that the *LTA4H* TT benefit was weaker and less prolonged in Indonesia Grade 2 than in Vietnam Grade 3 (compare Figure 1G to 1D). Non-TT patients had similar mortality in Indonesia Grade 2 and Vietnam Grade 3 (compare Figure 1G to 1D).

Thus, Bayesian analysis revealed an *LTA4H* TT survival association in Indonesia as well. The association being only in Grade 2 and not in Grade 3 patients suggested an upper limit of disease severity for its efficacy.

### Grade for grade mortality rate differences may explain the difference in grade-specific LTA4H TT effects in Vietnam and Indonesia

Why might *LTA4H* effects stop after Grade 2 in Indonesia? A closer comparison of the overall survival between the two sites suggested that there were mortality differences between the two cohorts for all grades combined and in grade for grade comparisons that were LTA4H-independent (Figure 1). For instance, Indonesia Grade 2 non-TT patients had a mortality risk similar to that of Vietnam Grade 3 non-TT patients (compare Figure 1D and 1G). We confirmed by Bayesian analysis that within each cohort, mortality risk increased with grade severity (Thuong et al., 2017; van Laarhoven et al., 2017) (Figure 2). From early on, Grade 1 survival was significantly greater than Grade 2, which was significantly greater than Grade 3 (Figure 2A and B). The hazard rate ratios highlighted that while the risk of increased mortality with higher grade was highest early, it was sustained long-term (Figure 2C and D). Importantly, both the survival and hazard rate analyses again pointed to an increased grade-for-grade risk of mortality in Indonesia over Vietnam.

**Figure 2.**
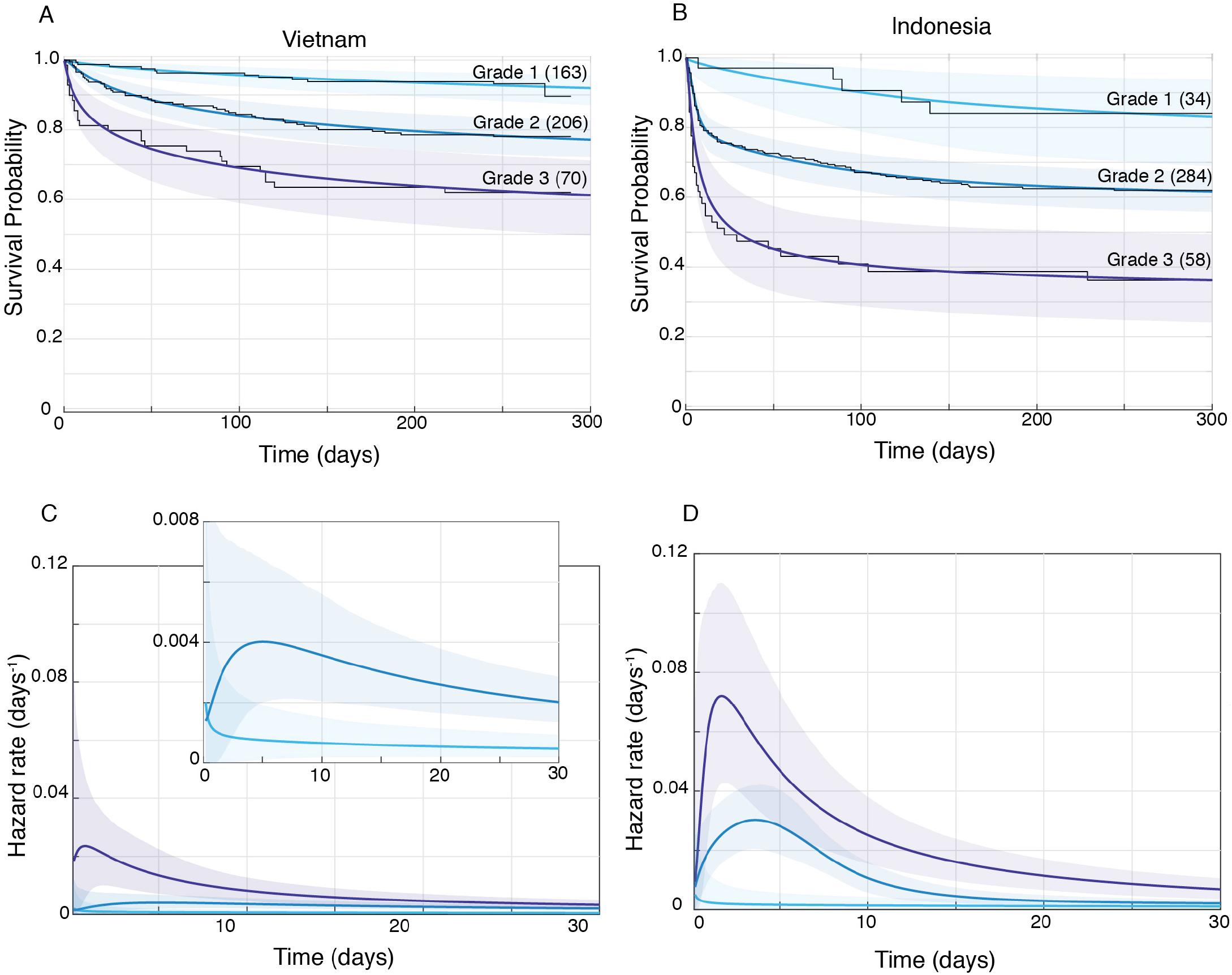
HIV-negative patient survival, stratified by grade. Mean posterior survival probability curves (coloured lines) overlaid by Kaplan-Meier survival plots (black lines) for Vietnam (A) and Indonesia (B), and mean posterior hazard rate curves for the first 30 days for Vietnam (C) and Indonesia (D). Shaded areas represent the 95% Bayesian confidence limits for posterior probability. The number of patients in each group at the starting time point are indicated in parentheses. (A) Vietnam Grade 1 over Grade 2 survival was significantly greater from day 7 onwards with maximum probability 0.999, survival gap 14%; Grade 2 over Grade 3 survival was significantly greater for from day 1 onwards with maximum probability 0.999, survival gap 16%. (B) Indonesia Grade 1 over Grade 2 survival was significantly greater from day 2 onwards with maximum probability 0.999, survival gap 21%; Grade 2 over Grade 3 survival was significantly greater from day 2 onwards with maximum probability 0.999, survival gap 25%. (C) Vietnam hazard rate ratio was >1 for both grade comparisons (inset magnifies Grade 1 and Grade 2 differences) with Grade 2 over 1 ratio peaking at 5.5 on day 7 and Grade 3 over 2 ratio peaking at 13.9 on day 1. (D) Indonesia Grade 2 over 1 hazard rate ratio was >1 up to day 215 and Grade 3 over 2 ratio was >1 throughout, peaking at 11.2 on day 1 for Grade 2 over 1, and at 2.3 on day 5 for Grades 3 over 2.

Next, we directly compared grade for grade mortality between the cohorts. Vietnam survival probability was higher in all grades, and significantly so for Grades 2 and 3 with survival gaps of 18% and 24%, respectively (Figure 3A, C and E). Similarly, hazard rates did not differ significantly between the cohorts in Grade 1 (Figure 3B), but there was a significant and substantial increase in early hazard rates for Indonesia Grades 2 and 3 (Figure 3D and 3F).

**Figure 3.**
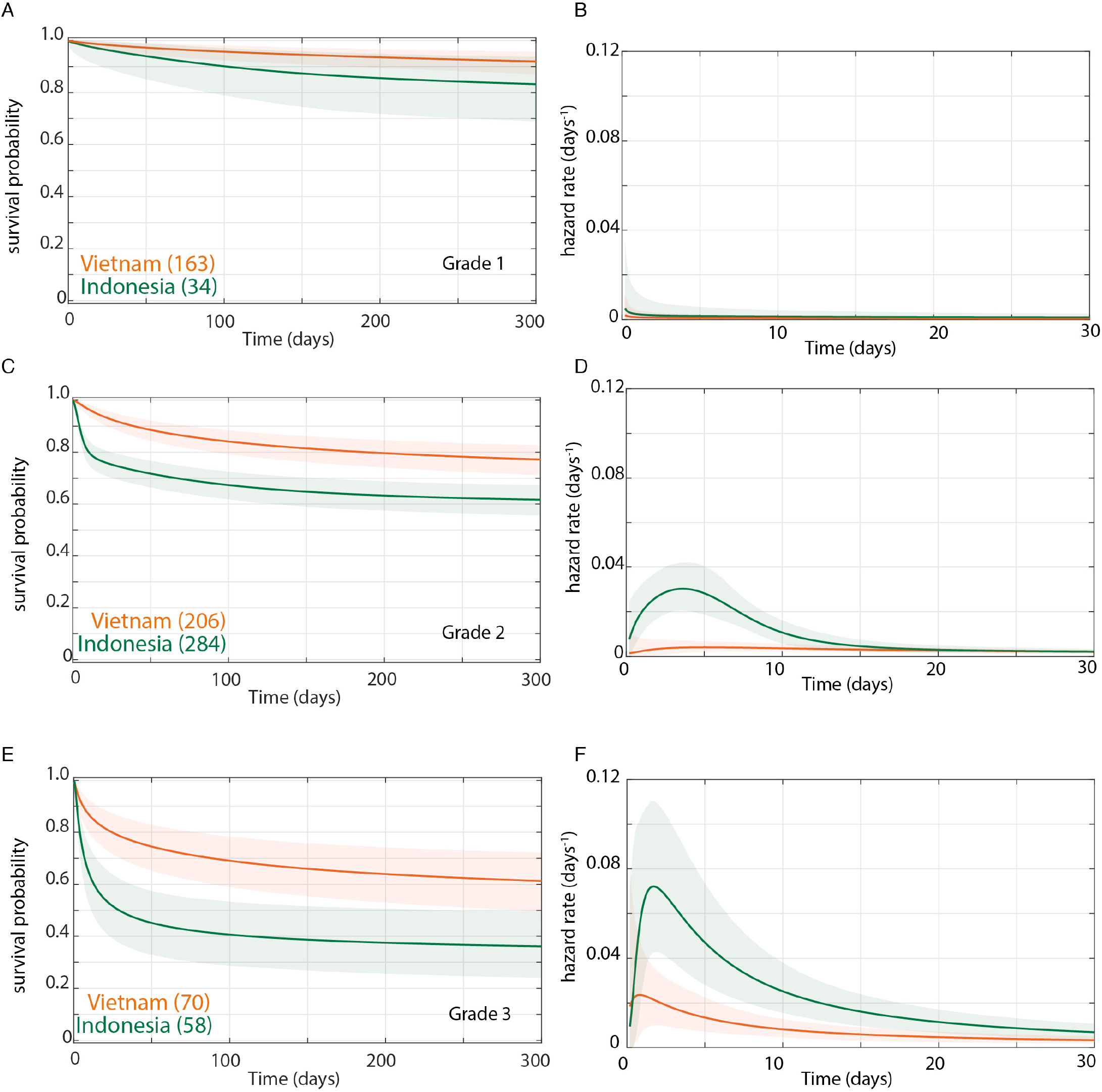
Grade stratified survival curves (A, C, E) and hazard rate curves for the first 30 days (B, D, F) for HIV-negative patients comparing Vietnam (orange lines) and Indonesia (green lines). The number of patients at the starting time point are indicated in parentheses. (A) Grade 1 survival did not differ significantly between Vietnam and Indonesia. (B) Indonesia hazard rate was ~2-fold higher than Vietnam in Grade 1 from day 0 to day 150, but the difference did not reach significance. (C) Grade 2 survival was significantly lower in Indonesia than Vietnam from day 1 onwards with maximum probability 0.999, survival gap 18%. (D) The hazard rate ratio in Grade 2 for Indonesia over Vietnam peaked at 8.4 on day 1 and remained >1 until day 180. (E) Grade 3 survival was significantly lower in Indonesia than Vietnam from day 2 onwards with maximum probability 0.999, survival gap 24%. (F) Grade 3 hazard rate ratio for Indonesia over Vietnam peaked at 3.6 on day 3 and remained >1 until day 165.

In sum, these analyses show that the inherent higher mortality associated with more severe disease on presentation was sharply accentuated in Indonesia. Indonesia grade 2 patients experienced similar mortality risk as Vietnam Grade 3 patients with the Indonesia Grade 3 patients experiencing far greater mortality. This higher mortality could potentially explain the finding that the *LTA4H* TT survival advantage did not extend to Indonesia Grade 3 patients. It may be that the TT genotype advantage, in response to corticosteroid treatment, is overridden by other factors that cause extreme mortality.

### LTA4H TT genotype is associated with increased survival in HIV-positive patients

HIV-positive TBM patients suffer ~ twice the mortality of their HIV-negative counterparts and the original RCT found that dexamethasone did not confer a significant survival benefit in HIV-positive patients (Stadelman et al., 2020, in press; Thwaites et al., 2004). The Vietnam study analyzed here also examined *LTA4H* genotype association with survival in dexamethasone-treated HIV-positive patients of similar ages and TBM grades to the HIV-negative cohort (Thuong et al., 2017) (Table 1). The TT association with survival was small, failing to reach statistical significance (Thuong et al., 2017). We noted that HIV-positive patients suffered the expected increase in overall mortality, twice that of HIV-negative patients (Table 1). Therefore, it was possible that in this cohort as in the Indonesia HIV-negative cohort, a weaker *LTA4H* TT survival effect that did not extend to Grade 3 was being masked in the overall cohort analysis. We first analyzed overall (*LTA4H*-independent) mortality differences in the HIV-positive patients. Like their HIV-negative counterparts, mortality rates were grade-dependent - Grade 3 > Grade 2 > Grade 1 (Figure 4A and B). A direct grade for grade comparison of the HIV-positive and HIV-negative patients showed that HIV-positive mortality was significantly higher in all grades (Figure 4C, E, G). While both groups had an early peak in hazard rate, this peak was higher in HIV-positive patients (Figure 4D, F, H). The HIV-positive hazard rate remained higher long-term, consistent with HIV-positive patients being broadly immunosuppressed. Thus, HIV-positive patients experienced an increased grade-for-grade mortality risk over HIV-negative patients.

**Figure 4.**
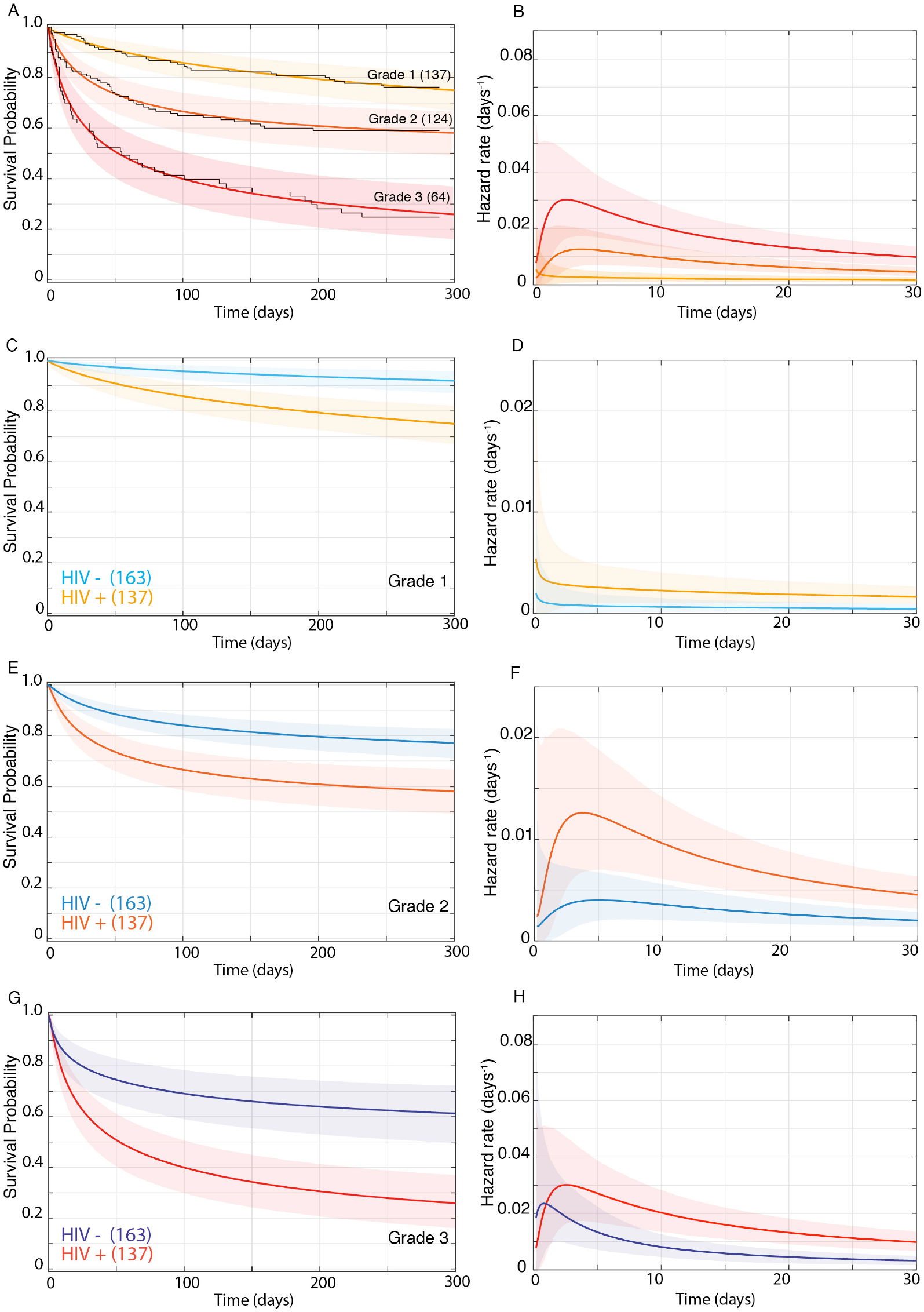
HIV-positive patient survival, stratified by grade. Mean posterior survival probability curves (A, C, E, G) and mean posterior hazard rate curves for the first 30 days (B, D, F, H). Shaded areas represent 95% Bayesian confidence limits. (A) Bayesian survival probability curves overlaid by Kaplan-Meier survival plots (black lines). Grade 2 survival was significantly greater than Grade 1 from day 5 onwards (maximum probability 0.99, survival gap 17%), and Grade 2 survival was significantly greater than 3 from day 3 onwards (maximum probability 0.99, survival gap 32%). (B) Hazard rate ratio was >1 from day 1 to day 176 for Grade 2 over Grade 1, peaking at 4.8 on day 5. For Grade 3 over 2 the hazard rate ratio was 3.3 at the start, fell to 2.1 on day 13, then rose steadily to 3.8 on day 270 and remained >1 throughout. (C-H) Grade-stratified nine-month survival of HIV-positive patients (orange lines) compared to their HIV-negative counterparts from Figure S2A (blue lines) (C, E, G).and their respective hazard rates (D, F, H). (C) Grade 1 HIV-positive survival was significantly lower than HIV-negative from day 11 onwards (maximum probability 0.999, survival gap 16%). (D) Grade 1 HIV-positive over HIV-negative hazard rate ratio was >3 throughout, peaking at 3.5 on day 65. (E) Grade 2 HIV-positive survival was significantly lower from day 4 onwards (maximum probability 0.999, survival gap 19%). (F). Grade 2 HIV-positive over negative hazard rate ratio was >1 throughout, peaking at 3.4 on day 2. (G) Grade 3 HIV-positive survival was significantly lower than HIV-negative from day 11 onwards (maximum probability 0.999, survival gap 35%). (H) Grade 3 HIV-positive over HIV-negative hazard rate ratio rose steadily to peak at 3.0 at 30 days and remained >1 throughout.

Bayesian analysis of *LTA4H* genotype effects in the overall HIV-positive cohort confirmed that TT patients did not survive significantly better than non-TT (Figure 5A). When we asked if the *LTA4H* effect was limited to less severe grades, we found that it was present in both Grades 1 and 3. *LTA4H* TT patients in both Grades 1 and 3 had a higher survival probability than non-TT (Figure 5C and G).

**Figure 5.**
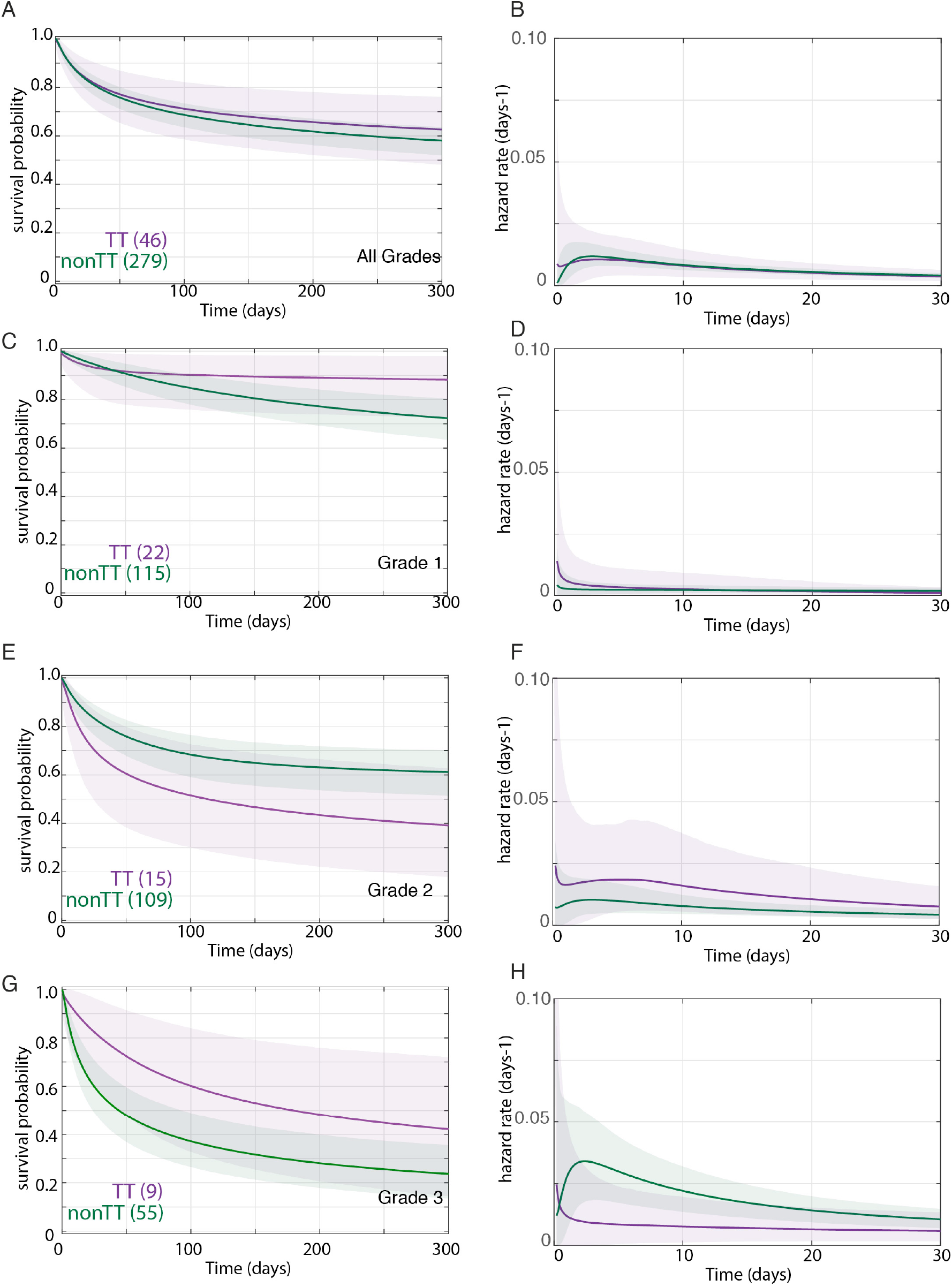
Effect of LTA4H rs17525495 genotype on HIV-positive patient survival probability for the nine-month observation period in Vietnam, overall (A) and stratified by grade (C, E, G), and hazard rates for the first 30 days, overall (B) and stratified by grade (D, F, H). Coloured lines represent mean posterior survival probability curves and mean posterior hazard rate curves. Shaded areas represent 95% Bayesian confidence limits. The number of patients at the starting time point are indicated in parentheses. (A, B) Overall TT and non-TT survival and hazard rates were not significantly different. (C, D) Grade 1 TT survival was higher from day 41, with the difference significant from day 241 onwards (maximum probability 0.95, survival gap 18%). The non-TT hazard rate was significantly lower than TT from day 48 onwards; the non-TT over TT hazard rate ratio was >1 from 14 days onwards, peaking at 8 at day 250. (E, F) Grade 2 non-TT survival was significantly higher than TT from day 252 onwards (maximum probability 0.95, survival gap 21% in favour of non-TT), but hazard rates did not differ significantly. (G, H) Grade 3 TT survival was significantly higher than non-TT from days 12-72 with maximum probability 0.97, survival gap 18%; hazard rate ratio for non-TT over TT peaked at 3.7 on day 3 and remained >1 throughout.

However, Grade 2 analysis gave an unexpected result. TT patients had lower survival probability than non-TT (Figure 5E). The patients had been admitted to one of two hospitals (Table S1 and Methods), and it appeared that this reversal in non-TT versus TT deaths was being driven substantially by three of the eight Grade 2 TT patients in Hospital 1 dying on day 7-8. In contrast, only 7 of the 68 non-TT patients admitted to Hospital 1 had died by day 8.

In sum, ignoring the likely spurious Grade 2 result, the *LTA4H* TT genotype was also associated with survival in HIV-positive patients, albeit with a weaker effect. Intriguingly, the effect persisted and indeed was strongest in Grade 3 despite this subgroup having a greater mortality risk than Indonesia Grade 3 HIV-negative patients (compare Figure 5G to Figure 1H). Rather, the increasing effect strength with grade mirrored the picture seen in Vietnam HIV-negative patients.

## DISCUSSION

The finding in two independent cohorts in Vietnam collected from 2001-2004 and 2011-2015 that a common functional human variant was associated with responsiveness to adjunctive glucocorticoid treatment in TBM represented an ideal example of pharmacogenomics, coming as it did from mechanistic understanding of the underlying reason (Sadée & Dai, 2005; Thuong et al., 2017; Tobin et al., 2012). The potential importance of these findings was heightened with subsequent pathway analyses in the zebrafish (where the *LTA4H* link to disease severity was discovered) identifying more specific, inexpensive, steroid-sparing drugs that might circumvent LTA4H-mediated pathology (Roca & Ramakrishnan, 2013; Roca, Whitworth, Redmond, Jones, & Ramakrishnan, 2019). It was disappointing and puzzling when a similar study of TBM patients in Indonesia failed to find a significant association (van Laarhoven et al., 2017). In a commentary published alongside the Indonesia and second Vietnam studies, Fava and Schurr suggested that background genetic differences between the cohorts might account for the lack of an *LTA4H* association (Fava & Schurr, 2017). However, noting that the Indonesia cohort presented with more severe disease and died earlier, they postulated that rather than invoking an unknown genetic phenomenon, it was more likely that the beneficial effects of dexamethasone for the *LTA4H* TT genotype were nullified in more severe disease. This concept of the disease being too severe for outcomes to be influenced by intervention has precedent. The beneficial effect of fluoroquinolones in TBM is present in Grade1 or 2 disease, but lost in Grade 3 disease (Thwaites et al., 2011). To test if increased grade severity on presentation in Indonesia was sufficient to account for the loss of the *LTA4H* TT survival effect, a grade for grade analysis within and between the two cohorts was necessary, while taking into account the temporal changes in mortality risk over the several months-long observation period. Moreover, both the magnitude and the time to mortality seemed to vary dramatically between the cohorts (Table 1). We realized that Bayesian methods were ideal for these complex and intrinsically multivariate comparisons and possibly the only path to a biologically and clinically relevant understanding.

When we analyzed the Vietnam cohort separated by grade severity, we saw that there was indeed a relationship between *LTA4H* effect and grade severity. However, it was opposite to what had been predicted (Fava & Schurr, 2017). The TT survival benefit became stronger not weaker with increasing grade. The analysis also suggested the reason for this. Patients with mild disease on presentation did well regardless of *LTA4H* genotype, so that the added benefit of the TT genotype was small. In Indonesia, the *LTA4H* TT effect was present in Grade 2 and completely absent in Grade 3, a finding that was perplexing until we analyzed the *LTA4H*-independent mortality risk of the two cohorts. The Indonesia cohort did not just contain a greater number of patients presenting at a more severe grade as had been noted earlier (Fava & Schurr, 2017), but patients in this cohort had substantially higher early mortality than their Vietnam counterparts even grade for grade (Table 1). Indonesia patients had nearly twice the mortality risk in Grade 2 and ~ 50% higher risk in Grade 3. Tellingly, Grade 2 Indonesia overall mortality risk was virtually identical to Grade 3 Vietnam (38% versus 37.9%), suggesting that this level of overall mortality represents the boundary of the beneficial effect of *LTA4H* TT. Thus, *LTA4H* TT efficacy was limited by other factors that cause mortality. These factors appear independent of severity grade on presentation, and if they exceed a threshold (represented by about ~ 40% mortality) then the beneficial effect of *LTA4H* TT is lost. Analysis of mortality risk enabled by temporal hazard rate analyses provided further insight. For all grades in both cohorts, the risk of death was greatest within days. So also was the excess grade for grade mortality risk in Indonesia. Likewise, the survival benefit of *LTA4H* TT also occurred early. Thus the *LTA4H* TT survival benefit was being nullified in Indonesia Grade 3 by unrelated factors increasing mortality risk in the same time period.

What might these factors be? Perhaps dysregulated inflammation had reached the point of no return in Indonesia Grade 3 TT patients in a manner not revealed by standard metrics of judging disease severity. If this were the case then glucocorticoids might no longer be beneficial. Since both TT and non-TT patients suffered identical excess mortality risk, its cause would be *LTA4H*-independent. Indonesia patients tended to be younger than Vietnam patients in all grades (Table 1), and perhaps were more prone to develop such a response. The more likely possibility is that better ancillary care was possible in Vietnam where all patients were enrolled into a clinical trial versus only 17% in Indonesia (Thuong et al., 2017; van Laarhoven et al., 2017). Optimized respiratory support, in particular, would be essential to keep patients alive through the early high risk stage in order allow for anti-inflammatory effects of glucocorticoids to benefit the TT patients.

Why was the *LTA4H* TT effect misunderstood to be limited to the least severe patients in both cohorts (Fava & Schurr, 2017)? Two reasons might explain this. First, it was not appreciated that apart from having fewer Grade 1 patients, Indonesia patients also suffered higher grade for grade mortality risk (Fava & Schurr, 2017). Second, because of a paucity of less severe patients, a subgroup analysis was performed in an attempt to tease out an effect in this group. Patients were divided into GCS 14-15 (less severe) versus < 13 (more severe), and a nonsignificant TT survival benefit was found in the GCS 14-15 group only, supporting the idea that TT effects, if any, were limited to the less severe group (Supplementary Figure 2A and B). However, our analysis now shows the two reasons why this grouping resulted in the *LTA4H* TT effect going unrecognized: 1) GCS 15 does not separate patients by whether or not they have neurological signs, so it includes all BMRC Grade 1 and some BMRC Grade 2 patients (Box 2); 2) the grouping used resulted in the Grade 2 patients being split so that some were grouped with Grade 3 and the remaining with Grade 1; 3) of the 11 TT patients in the GCS 14-15 grouping only 1 TT patient was in Grade 1, with the remaining 10 in Grade 2, so the effect seen in their GCS 14-15 was entirely from Grade 2 patients (compare Figure S2B to S2C). In any case this subgroup TT effect would have been impossible to detect with the frequentist analyses used.

Grade-specific Bayesian analysis of the HIV-positive TBM patients in the Vietnam cohort reveals a TT benefit in these patients as well. Grade-specific Bayesian analysis showed an overall salutary effect on survival in Grades 1 and 3. As discussed in the Results section, we suspect that the likely spurious reversal of the TT and non-TT survival benefit in Hospital 1 Grade 2 prevented finding a TT benefit on overall survival. It is difficult to formulate a pathophysiological mechanism that would explain this reversal in the intermediate grade exclusively in HIV-positive patients, while still maintaining the TT survival advantage in both lower and higher severity. We note that, having reported on a long series of Bayesian significant results where each result gives a 0.95 posterior probability that a particular hypothesis is true, we might expect on average one in twenty of those hypotheses to be false, despite the result (Box 1). This particular result (with a posterior probability of 0.962) seems most likely to be one such, given that we have ~ 30 significant results in this analysis, and that the data for this group may be skewed by a small cluster of deaths at a single time point in one hospital. Therefore, the TT effect is most likely to occur across all grades, and by extension glucocorticoids are likely to provide benefit to TT individuals in all grades, albeit a weaker effect than their HIV-negative counterparts. That the TT effect would be present in HIV-positive Grade 3 patients might seem surprising at first blush because this group has an even higher mortality risk than Indonesia Grade 3 HIV-negative patients (75.2% versus 63.8%) (Table 1). The time of mortality in these groups provides a likely explanation. The median time to mortality was four times longer (37 days for Vietnam HIV-positive vs. 8 days for Indonesia). These analyses suggest that HIV-positive Grade 3 patients have a more prolonged risk of mortality for reasons that are independent of corticosteroid treatment and *LTA4H* genotype. Thus, the use of Bayesian methods has identified HIV-negative subgroups of TT patients that gain the most survival benefit from dexamethasone treatment.

HIV-positive TT patients also have an early survival benefit albeit weaker than in their HIV-negative counterparts. This may reflect a distinct pathophysiology from early on. This idea is supported by a subgroup analysis in the original Vietnam study which found that *LTA4H* TT did not associate with survival in severely immunocompromised individuals with a CD4 T lymphocyte count <150. Rather it appeared to be limited to those with CD4 >150, although the small number of patients in this group precluded finding significance. Together, our findings set the stage for using Bayesian methods for subgroup analyses of two ongoing trials where 1) the survival of HIV-negative CC+CT patients randomized to getting dexamethasone or placebo is being compared to that of TT patients who are all getting dexamethasone (Donovan, Phu, Thao, et al., 2018), and 2) the benefit of dexamethasone is being examined for HIV-positive patients of all three genotypes by randomizing them to get dexamethasone or not (Donovan, Phu, Mai, et al., 2018).

Finally, in addition to providing guidance for TBM pharmacogenomic approaches, we hope that our analyses highlight the unique value of Bayesian methods for providing guidance for other complex diseases with difficult treatment decisions. The vital importance of defining the patient populations and subgroups which will benefit the most from specialised interventions and treatments is increasingly appreciated (Sadée & Dai, 2005). This is not only to target such treatments to those who will benefit, but to avoid their adverse effects in those individuals who have little chance of experiencing a clinically relevant benefit from them. Our finding that the *LTA4H* TT genotype’s salutary role is incumbent on the optimization of other factors that maximize patient survival has broad implications for pharmacogenomic approaches.

## Acknowledgments

We thank R. Troll for evaluating the two published cohort studies, realizing that Bayesian analysis could provide answers and initiating the collaboration with the Bayesian statistician R.S. This work was supported by the NIH (NIAID ULTIMATE project 1R01AI145781-01) to A.L. and R.C., by the Wellcome Trust to N.T., T.T. and L.R. and by an NIH MERIT award to L.R.

## Author Contributions

L.W. participated in prior selection, analyzed and interpreted data, wrote paper; J.C wrote code, reviewed Supplementary Methods section; A.L., N.T., S.D, B.A, A.G.,R.C. provided raw data, discussed results, reviewed paper; G.T. participated in prior selection, provided raw data, discussed results, guided writing of, and edited, paper; M.T., P.E. participated in prior selection, reviewed data analysis, provided critical input, edited paper; R.S. checked code and added output, plotted data, performed housekeeping data management functions, analysed and interpreted data, wrote paper; L.R. conceived and oversaw project, participated in prior selection, guided data analysis, interpreted data, wrote paper.

## Box 1.

### Contrast of “95% significant” in Bayesian and frequentist paradigms (MacKay, 2003)

Bayes: “A is significantly greater than B” = Posterior probability that A greater than B is at least 0.95.

Frequentist: “A is significantly greater than B” = For any circumstance where A is at most B, the probability of getting data in this critical region, as we did, was at most 0.05.

Therefore:

1. We expect 1 in 20 of Bayesianly (95%) significant results to be truly negative and therefore false positives;
2. We expect up to 1 in 20 of truly negative results to be frequentistly significant (at the 95% level) and therefore false positives.

Therefore (assuming all positives are at 95% level):

- In the frequentist paradigm, the expected number of false positive results is proportional to the number of comparisons done on true negatives;
- In the Bayesian paradigm, the expected number of false positive results is proportional to the number of apparent positive results, and unaffected by any vast number of accompanying apparent negative results.

## Box 2.

### TBM disease severity classification

*Glasgow Coma Score (GCS)* A general measure of consciousness used for a wide range of neurological deficits, particularly brain trauma, by scoring eye opening and verbal and motor responses to stimuli, to assign a numerical value from 3-15 corresponding to decreasing severity, where 3 corresponds to completely unresponsive, deep coma and 15 to fully conscious (Teasdale & Jennett, 1974).

*Modified British Medical Research Council (BMRC) TBM Grade* A classification scheme specifically tailored to assess TBM severity. It is derived from the GCS, and additionally incorporates the presence of focal neurological signs. The TBM grade is scored between 1-3 corresponding to increasing severity, converse to the GCS classification.

*GCS and BMRC Grade relationship* BMRC Grade 1 - GCS=15 with no focal neurological signs; BMRC Grade 2 - GCS=11-14, or GCS=15 with focal neurological signs, BMRC Grade 3 - GCS<10 (Heemskerk et al., 2011).

## Box 3

### Definitions and usages

Definitions

- *Posterior probability* - the probability after seeing the data
- *Mean posterior survival probability at time T* - the expectation after seeing the data of the fraction of patients that will still be alive at time T
- *Hazard rate* - the fraction of those still surviving that will die per unit time. A high hazard rate at a particular time indicates that patients are at high risk of dying at that time
- *Mean posterior hazard rate at time T* - the expectation after seeing the data of the hazard rate at time T

### Abbreviated and example usages

*Onwards* - for the rest of the 270-day observation period

*Throughout* - for the entire 270-day observation period

*“A was significantly greater than B at time T”* - the posterior probability that A was greater than B, at time T, was at least 0.95

*“A was not significantly different from B”* - the posterior probability that A was greater than B was between 0.05 and 0.95 throughout

*“Group A survival was 30% greater than group B”* - the mean posterior survival probability at 270 days, p_A_, was 30% absolute greater than the corresponding probability p_B_ for group B. (“*absolute*” here meaning that p_A_= p_B_ + 0.3, and not that p_A_ = p_B_ × 1.3)

*“Probability that group A survival was better than group B at time T was 0.97”* - the posterior probability that group A survival probability at time T was greater than group B survival probability at time T was 0.97 (Note that this is not a reference to the mean posterior probability.)

*“The hazard rate ratio for group A over group B peaked at X at time T and remained greater than Y throughout”* or *“Group A had an X-fold higher relative risk of death at time T which remained greater than Y throughout”* - the mean posterior hazard rate for group A, divided by that for group B, peaked at a value of X at time T and remained greater than Y at all times up to 270 days.

*“The probability that hazard rate for group A was greater than that of group B was 0.97 at time T”* - the posterior probability was 0.97 that the hazard rate for group A at time T was greater than that for group B at time T (Note that this is not a reference to the mean posterior hazard rate.)

“*The hazard rate for group A was greater than that for group B at time T*” - the mean posterior hazard rate for group A was greater than that for group B at time T.

“*survival gap”* - the difference in mean posterior survival probability between the two groups being considered at 270 days

---

**Table S1.** Characteristics of the HIV-positive Vietnam cohort separated by hospital.

| **Hospital** | **1** | **2** |
| --- | --- | --- |
| **Total** | 187 | 138 |
| **Age in years**<br>*median (range)* | 33 (18-53) | 33 (22-63) |
| **Glasgow Coma Score**<br>*median (range)* | 15 (5-15) | 14 (3-15) |
| **BMRC TBM grade**<br>*no. (% of total)* |  |  |
| **1** | 90 (48.1) | 47 (34.1) |
| **2** | 68 (36.4) | 56 (40.6) |
| **3** | 29 (15.5) | 35 (25.4) |
| **Overall mortality**<br>*no. (%)* | 76 (41.2) | 53 (39.3) |
| **Time to median mortality**<br>*(days)* | 37 | 34 |
| **Mortality by BMRC TBM grade**<br>*No. dead (% of grade)* |  |  |
| **1** | 20 (22.8) | 12 (26.0) |
| **2** | 33 (49.2) | 17 (30.8) |
| **3** | 23 (79.3) | 24 (71.8) |

**Figure S1.**
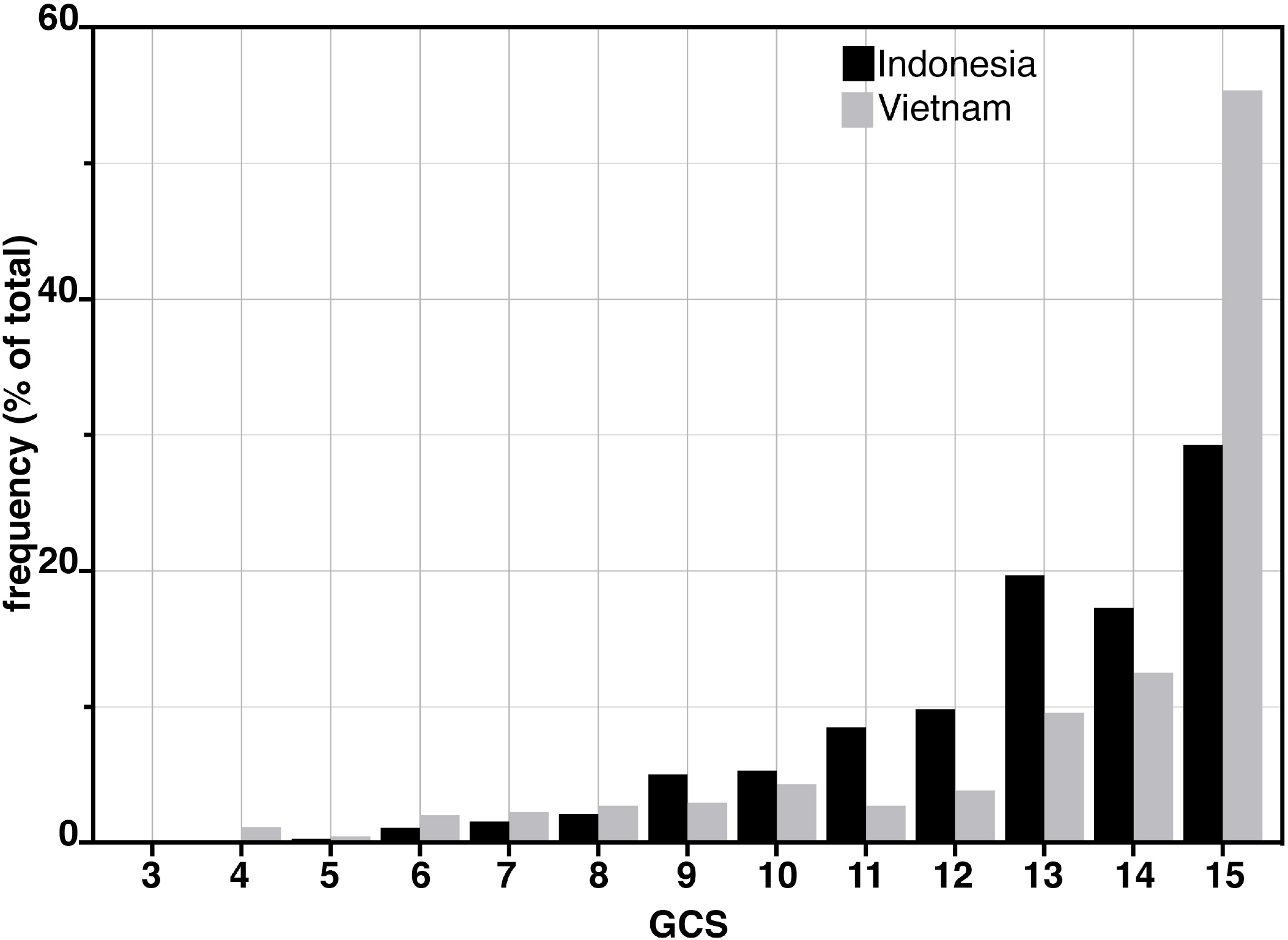
Glasgow Coma Scores (GCS) for Vietnam and Indonesia HIV-negative patients. Frequency of GCS values indicated on the Y-axis as a percentage of the total cohort (n=376 Indonesia, n=439 Vietnam).

**Figure S2.**
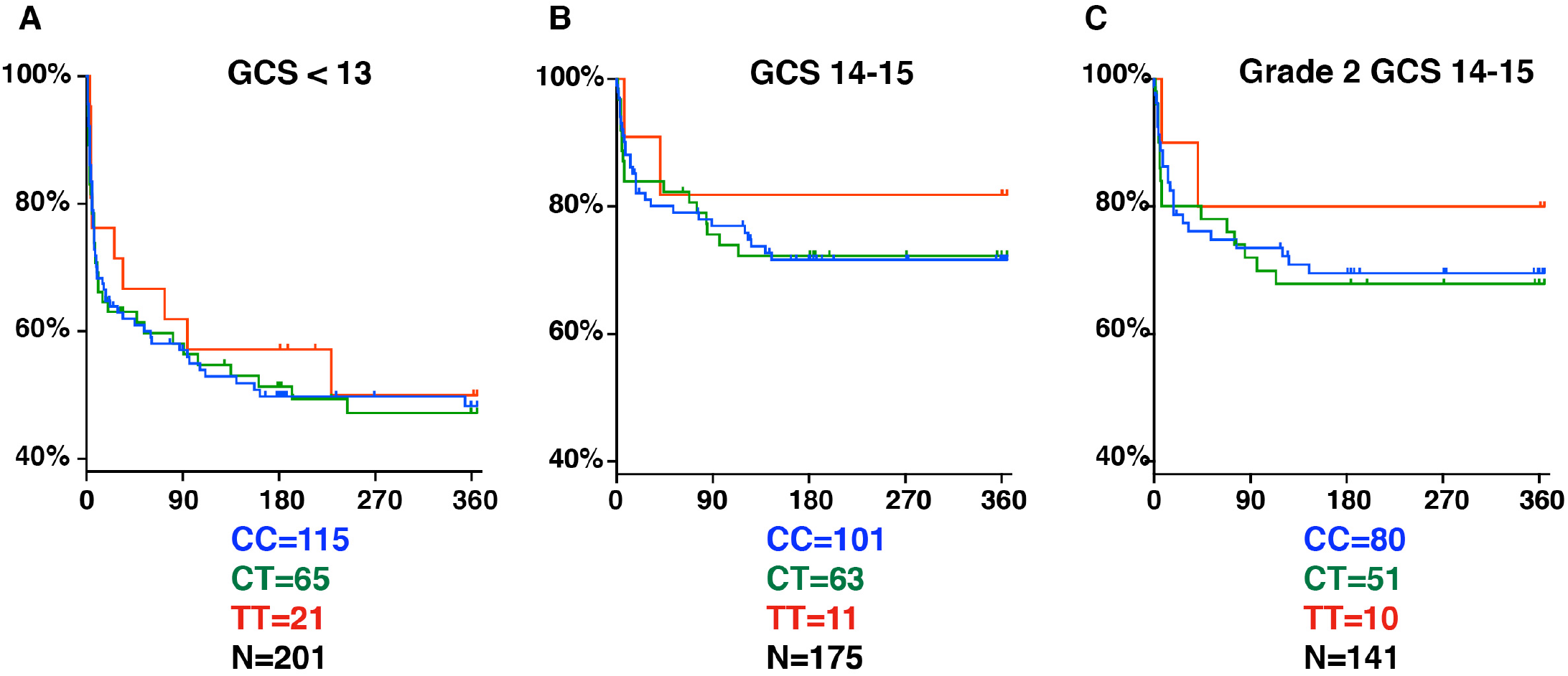
LTA4H rs17525495 genotype as predictor for 365-day mortality in HIV-negative TBM patients in Indonesia. Panels A and B are comparable to Figure S2B of van Laarhoven et al. 2017. (A) All patients with GCS<13. (B) Patients with GCS=14 or 15 (note that 1 patient with TT genotype, GCS=15 was censored on day 30 in the original data set but later found to have died on day 41). (C) The subset of patients from (B) with GCS 14 or 15 excluding those in BMRC Grade 1 (GCS 15 without neurological signs).

## Supplementary Methods

## 1 Overview

We adopt the Bayesian paradigm. Accordingly we define below a probabilistic generative model for patient lifetime *x* given model parameters *θ* and a suitable prior distribution on *θ*. We assume that, when observed, such lifetime data may be censored at known times *t* that are independent of the underlying lifetimes and the model parameters, so that for each patient we observe either a time of death *x* or a time of censoring *t* at which the patient was alive.

We then collect a data set of values 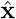 of *x* or *t* for a set of patients, and apply Bayes’ theorem to deduce the posterior distribution 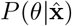 of the model parameters.

Since this distribution is hard to visualise, we draw samples of *θ* from it using Markov chain Monte Carlo (MCMC) methods described below. For each such sample of *θ* we calculate the distribution of lifetime *x* given *θ* and hence the survival probability *q*(*t*|*θ*) and hazard rate *h*(*t*|*θ*) functions against patient time since admission, using the following relationships:

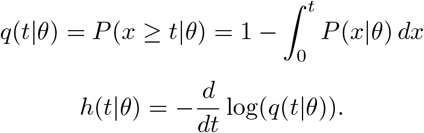

An example resulting set of survival probability curves is shown in Figure 1, with the corresponding hazard rate curves being shown in Figure 2.

**Figure 1:**
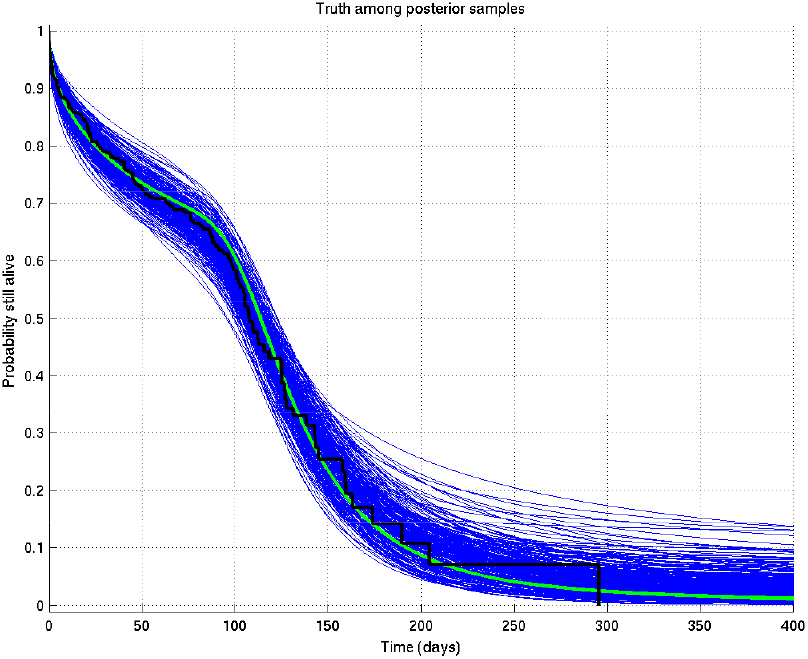
Output of test run using synthetic data for which the right answer is known. The true survival probability curve is shown in green, with the Kaplan-Meier plot for the generated data in black. In blue are shown many samples from the posterior distribution on the survival probability curve, calculated from 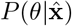, which indicate the uncertainty in the inferred distribution. The synthetic dataset comprised 300 hypothetical patients of whom the time of death of 153 was censored.

**Figure 2:**
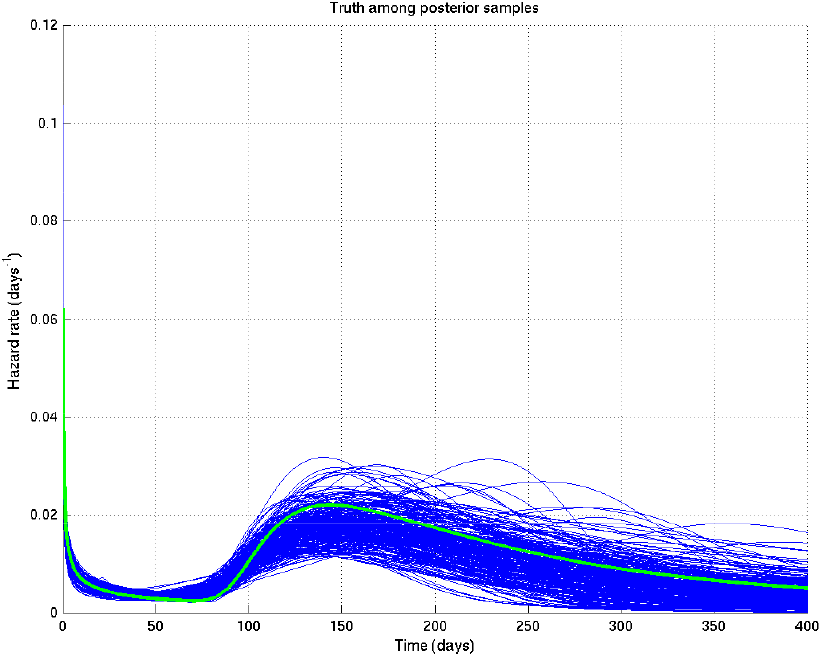
Output of test run using synthetic data for which the right answer is known. The true hazard rate curve is shown in green. In blue are shown many samples from the posterior distribution on the hazard rate curve, calculated from 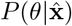, which indicate the uncertainty in the inferred distribution. The synthetic dataset comprised 300 hypothetical patients of whom the time of death of 153 was censored.

Now, given a set of such curves, we can calculate the mean posterior survival probability (resp. hazard rate) at each time point and plot the posterior mean survival probability (resp. hazard rate) against time. Similarly we can find the 2.5% and 97.5% centiles. Examples of the posterior mean and centile curves for survival probability corresponding to Figure 1 are shown in Figure 3.

**Figure 3:**
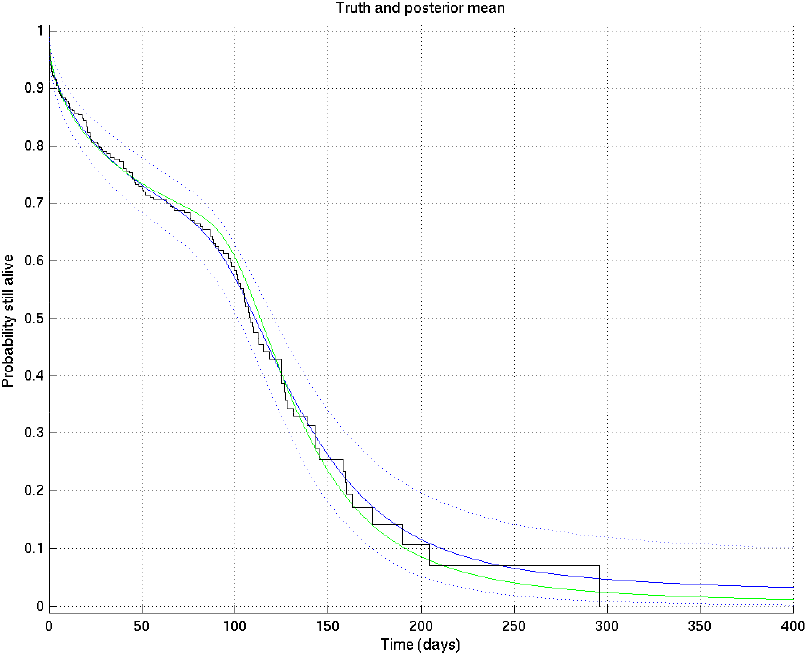
Output of test run using synthetic data for which the right answer is known. The true survival probability curve is shown in green, with the Kaplan-Meier plot corresponding to the generated data in black. In blue is the posterior mean survival probability against time, calculated from 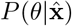, and in dotted lines the 2.5% and 97.5% centiles, which indicate the uncertainty in the inferred distribution. The synthetic dataset comprised 300 hypothetical patients of whom the time of death of 153 was censored.

Further, given two such sets of survival probability or hazard rate curves (such as the survival probability curves shown in Figure 4 and zoomed in in Figure 5), we can also calculate at each time point the probability that a value taken at that time from a random curve from set 1 is greater than a similar value taken from a random curve from set 2. As we can see from these two figures, this probability is higher at 10 days than at 1 day or 100 days. These probabilities, at each time point, can then be plotted giving in this example Figure 6, showing the probability that underlying survival in group 1 is greater than that in group 2 at each time point.

**Figure 4:**
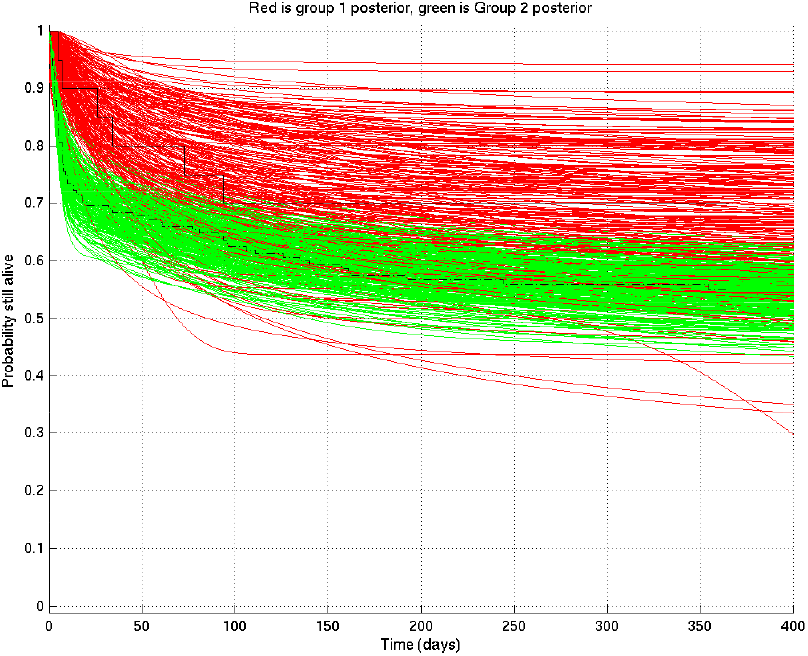
Comparison of two runs generated independently from two subsets of patients, subset 1 (red) consisting of 20 patients and subset 2 (green) consisting of 182 patients. The Kaplan-Meier plot for subset 1 is in solid black and that for subset 2 in dot-dashed black. Since there were many more patients in subset 2 than in subset 1, we expect greater variance in the inferred survival probabilities for subset 1 than for subset 2.

**Figure 5:**
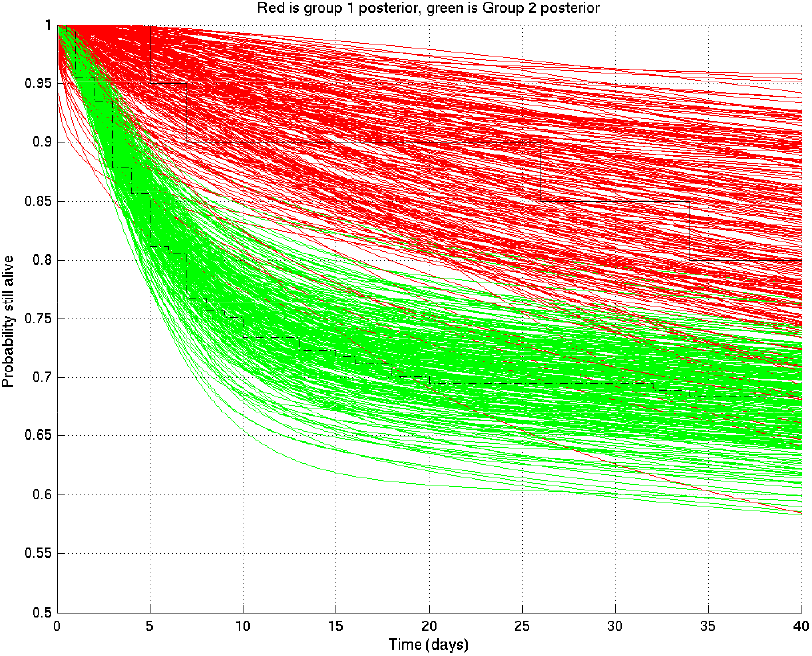
As for Figure 4 but zoomed in to the top left hand corner, showing greater separation and less overlap of the red and green curves at 10 days than at 1 day or 100 days.

**Figure 6:**
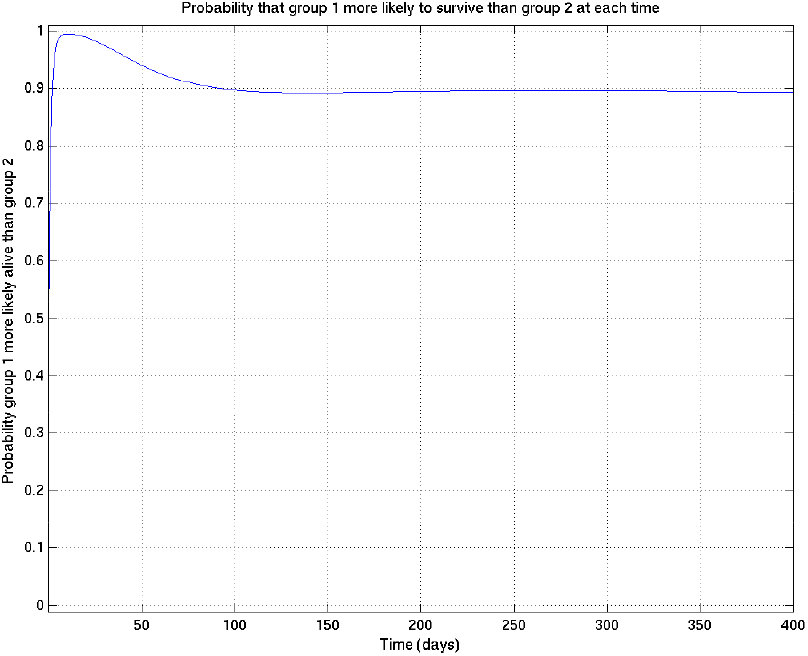
Corresponding to Figures 4 and 5 this shows the probability that survival for subset 1 is greater than that for subset 2 at each time point.

In order to carry out this process, we need to specifically define the lifetime distribution model *P* (*x*|*θ*), and we now turn to this.

## 2 Lifetime model

We suppose that there exist an unknown number *J* of different mechanisms causing death, and that each such mode of death has a different lifetime distribution.

### 2.1 Combination of different modes of death

Let *x* denote a lifetime, i.e. the time until a patient dies. Let *j* ∈ {1, 2,…, *J*} denote a particular mechanism (or mode) of death. Let *x_j_* denote the time at which mode *j* would kill the patient; we set *x_j_* = ∞ to denote the possibility that that mode would never have killed the patient.

Then the patient’s time of death is given by

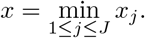

In particular *x* = ∞ denotes the situation that the patient never dies (unlikely as this is).

### 2.2 Model of a single mode of death

We now drop the subscripts *j*, but assume that this subsection will be repeated *J* times with the subscript *j*s added to every random variable, with each of the repetitions being independent as far as the model is concerned before being conditioned on observed data. When later we want to refer to the complete set of *J* values of e.g. *p*, we will use bold face, e.g. **p** = (*p*_1_, *p*_2_,…, *p_J_*).

Thus we will set *P*(*x*|*p, k, m, r*), i.e. *P*(*x_j_*|*p_j_, k_j_, m_j_, r_j_*), to be such that with probability *p*, *x^k^* is Gamma distributed with parameters *m′* = *m* and *r′* = *mr^k^*, and otherwise *x* = ∞. Thus we have

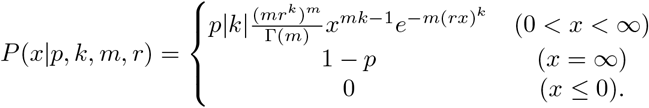

Here *p* ∈ [0, 1], *k* ∈ ℝ \ {0}, *m, r* > 0.

Note that we have here a distribution which has both a discrete and a continuous part, so that *P* (*x*|…) is used as notation both for a probability and for a probability density: in other words, we have a continuous distribution for finite positive *x*, given by a density function, whose interpretation is that its integral from *x*_1_ to *x*_2_ is the probability that *x*_1_ < *x* < *x*_2_; but as we have a non-zero probability 1 − *p* that *x* = ∞, the integral from 0 (inclusive) to ∞ (exclusive) of the density given by the first line of the above formula for *P* (*x*|*p, k, m, r*) must be *p*. On the other hand we have a discrete distribution for *x* = ∞, and 1 − *p* is a probability, not a density.

By way of very approximate intuition: *p* is the probability that a particular mode of death would kill the patient at a finite time; *r* is the reciprocal of the overall timescale to deaths of those patients who die; *m* governs how variable those times of death are – the smaller *m* is, the more variable are the times of death; and the sign of *k* plays a part in determining whether the hazard rate for this mode of death is increasing or decreasing, while the magnitude of *k* governs how abruptly the spread of death time is cut off in the less spread out direction. Specifically, *k* = 0 makes no sense, as then we would have *x^k^* = 1 for all *x*, and an invalid distribution would result (so it should not be a surprise that the prior on *k* is bimodal with zero density at zero).

### 2.3 Priors on the parameters

We specify the priors on the parameters in two stages. First, we specify their general form, and second we choose specific values for the hyperparameters that then specify a unique prior.

#### 2.3.1 General form of the priors

The total number *J* of modes is itself to be considered a random variable, on which we put the prior

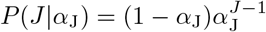

for *J* ∈ ℕ* = {1, 2,…} and for some fixed *α*_J_ ∈ [0, 1).

The prior for the parameters *p, m, r, k* of each mode of death are taken to be independent, and as follows. We take the prior on *p* to be Beta, with positive real parameters *α*_p_, *β*_p_ > 0, so that

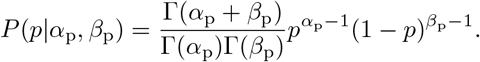

We take the prior on *r* to be Gamma, with parameters *m*_r_, *r*_r_ > 0, so that

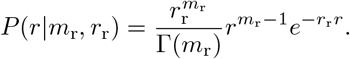

We take the prior on each of the parameters *k* and *m* to be the conjugate prior on each with respect to this parameterisation. Thus for positive real parameters *a*_m_, *b*_m_ we have

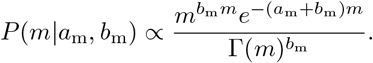

Similarly for parameters *N*_k_ ∈ ℕ, *a*_k_ ∈ ℝ_+_ and 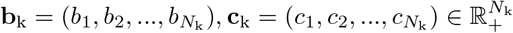 we have

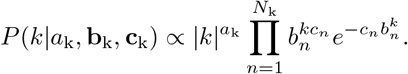

#### 2.3.2 Specific values of the hyperparameters and the resulting priors

Specific values were chosen for the hyperparameters by varying them and showing those users overseeing the analysis (namely LW, MT, PE, LR, of whom PE and LR are infectious diseases clinicians) the resulting distributions of samples of survival curves and hazard rates, then letting them choose the most appropriate prior given their background experience. The prior chosen was intentionally uninformative and very wide, while still being centred on the clinician-expected survival curves and hazard rates.

The specific parameter values chosen were as follows:

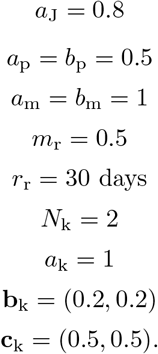

These result in the following depicted distributions for *J, p_j_, m_j_, r_j_, k_j_*, and hence for the depicted samples from the distributions for survival probability and hazard rate against time as well as the mean and 2.5% and 97.5% centiles for the last two: see Figures 7 to 15.

**Figure 7:**
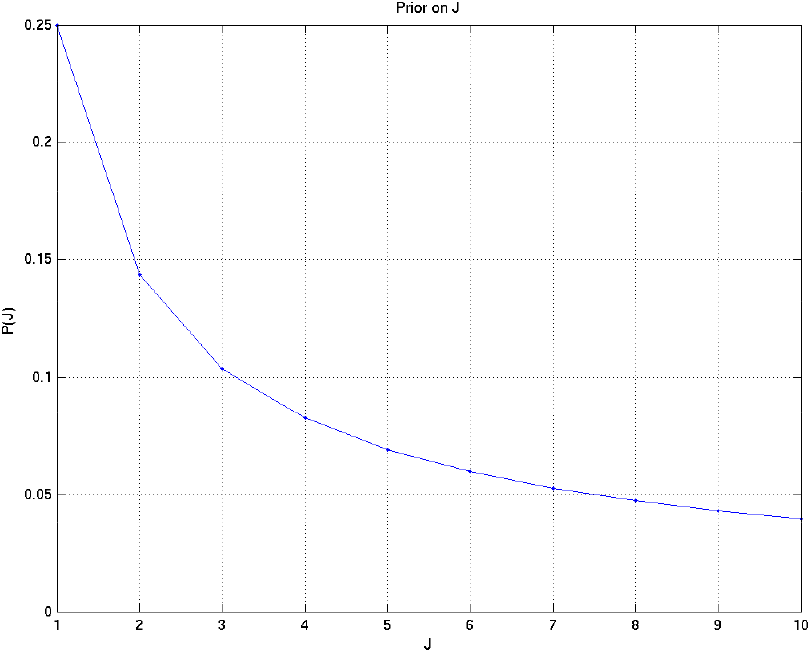
Prior on *J*, the number of different modes of death.

**Figure 8:**
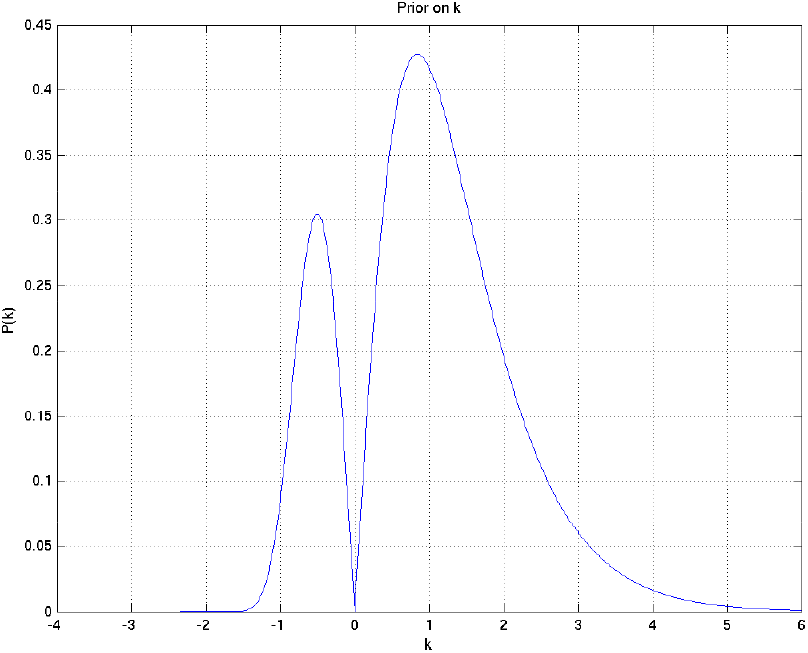
Prior on *k*.

**Figure 9:**
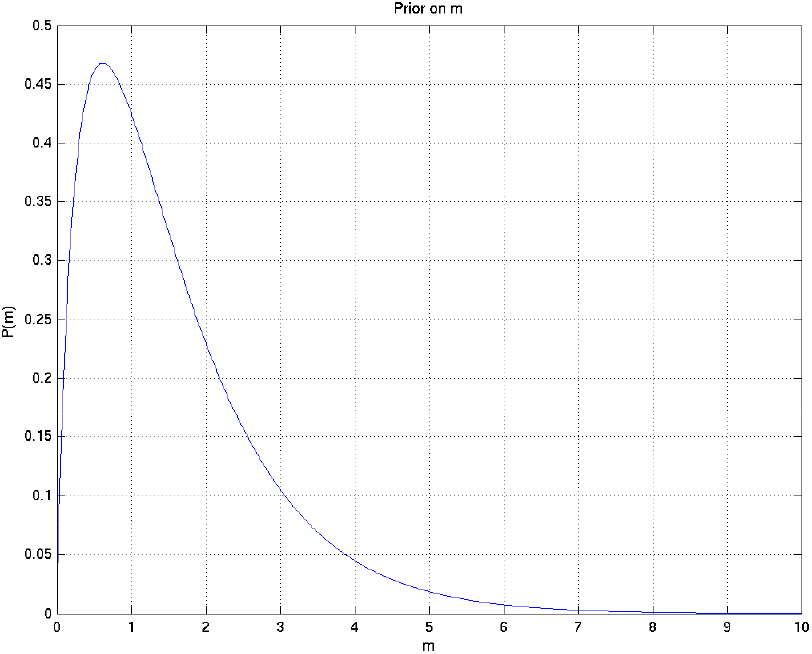
Prior on *m*.

**Figure 10:**
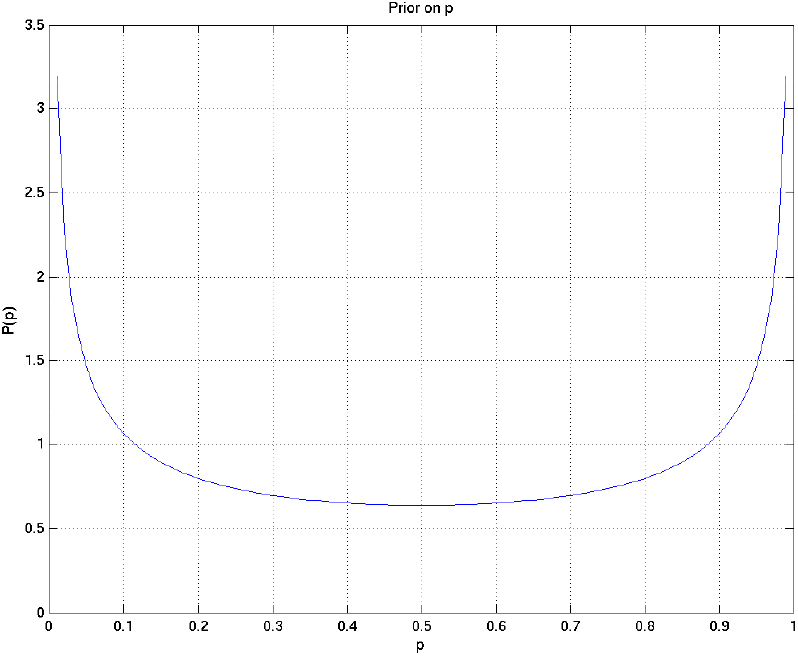
Prior on *p*.

**Figure 11:**
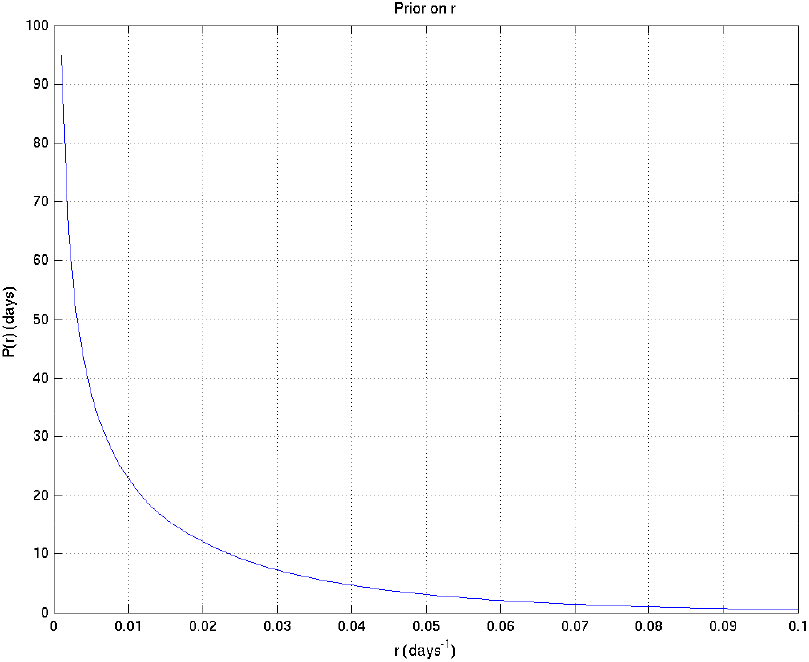
Prior on *r*.

**Figure 12:**
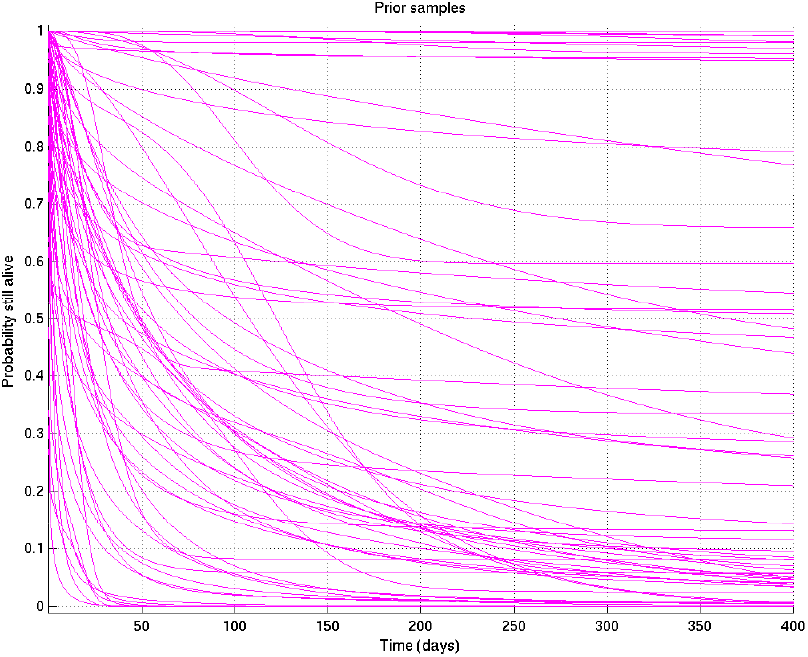
Samples from resulting prior on survival probability against time.

**Figure 13:**
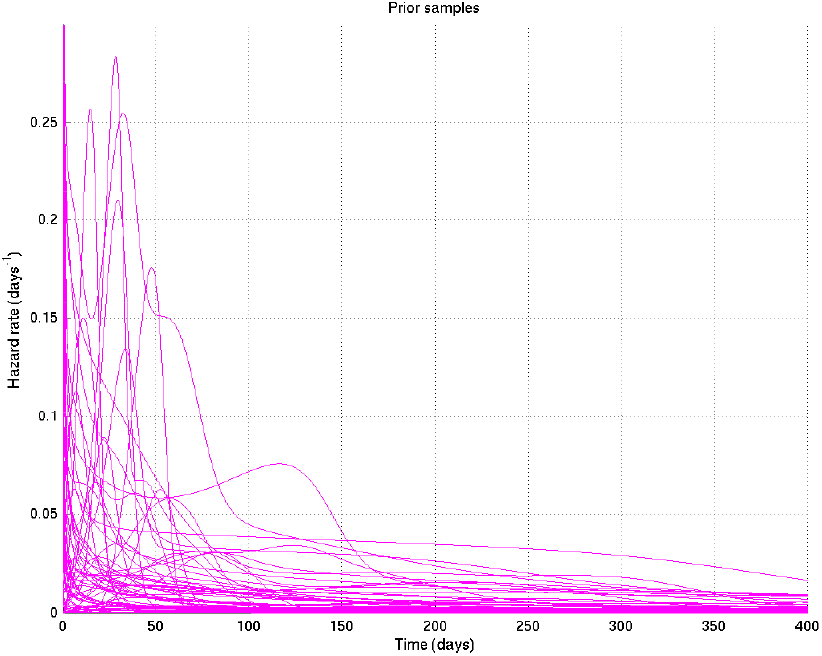
Samples from resulting prior on hazard rate against time.

**Figure 14:**
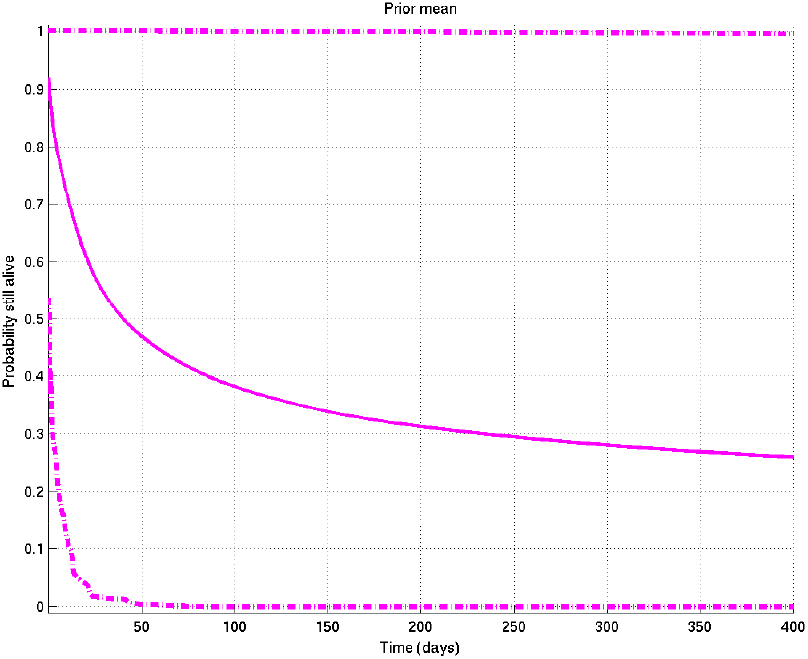
Mean and 2.5% and 97.5% centiles of prior on survival probability against time.

**Figure 15:**
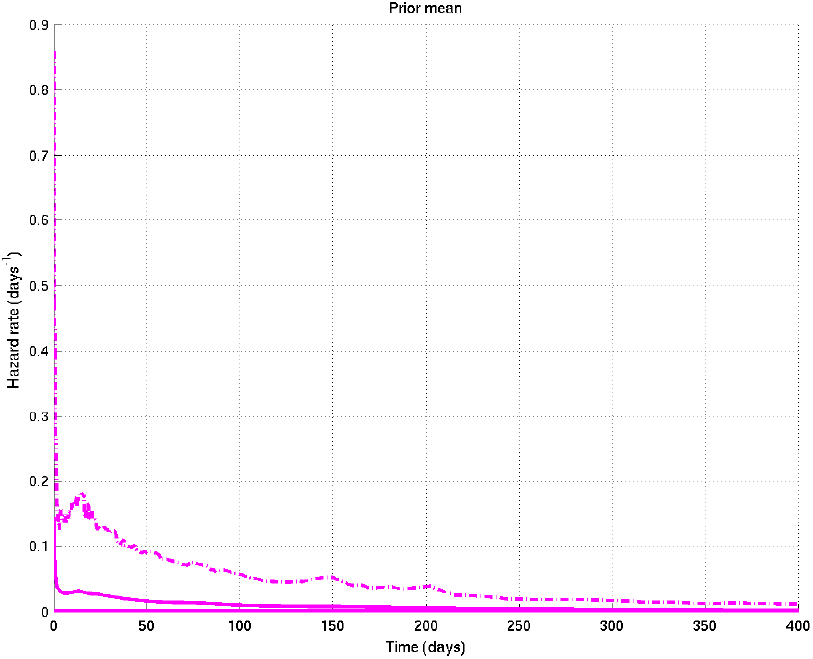
Mean and 2.5% and 97.5% centiles of prior on hazard rate against time.

### 2.4 Summary

Thus the parameters *θ* consist of *θ* = (*J,* **p**, **m**, **r**, **k**), and the prior

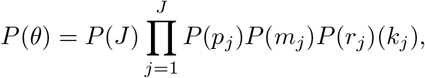

a combination of densities of different dimensionalities. This can be regarded a mixture of models, one for each possible value of *J*, weighted by the prior on *J*: the model for *J* = 1 has only one possible mode of death, while that for *J* = 2 has two possible modes of death, which are independent, and so on for higher values of *J*.

In particular, when comparing two subsets A and B of patients, we use the same prior to infer the posterior distribution of *θ* given 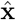 for each subset of the patients independently. Thus *a priori* the probability at each time *t* that *q*_A_(*t*) > *q*_B_(*t*) is 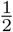, the probability that *q*_A_(*t*) < *q*_B_(*t*) is 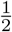, and the probability that *q*_A_(*t*) = *q*_B_(*t*) is zero (though it has non-zero probability density). In other words we are asking how sure we are that *q*_A_(*t*) > *q*_B_(*t*) given the data, reckoning the alternative to be that *q*_A_(*t*) < *q*_B_(*t*), and taking for granted that the probability that the two are *exactly* the same is zero.

### 2.5 Rationale

The basic Gamma model is a frequently used model of a failure mode that goes through several stages of failing, each with an exponentially distributed lifetime, before final failure occurs. The additional effect of the parameter *k* is to incorporate Weibull-type failure time models, allowing for both increasing and decreasing hazard rates. The parameter *p* allows for the possibility that some modes of dying may not be relevant for all patients, while the combination of the *J* submodels allows for a number of different types of mechanisms of death to be relevant.

In particular, we specifically used independent priors on the parameters of the subsets being compared because:

- When offered the choice the users (namely LW, MT, PE, LR, GT of whom PE, LR, and GT are infectious diseases clinicians with GT having extensive experience with TB meningitis patients) indicated unanimously that this correctly represented their prior beliefs;
- Because we restrict our use of posterior probabilities to those of the form *P* (*U > V*| data) rather than *P* (*U > V* +*α*| data) for some *α* > 0, and because the additional effect of any non-independent factors in the prior would be symmetric either side of the difference between the two subsets being zero, the effect of any non-independent factors would be expected to be very small;
- If they did not believe the priors on the relevant disjoint subsets were independent, the clinicians involved would find it very hard to specify exactly how similar each pair of subsets being compared should be expected to be.

## 3 MCMC methodology

We introduce additional variables *j_i_* for each patient *i* which indicate whether the time of death was censored (value 0) or was caused by a particular mode *j* of death (value *j* ≠ 0 unknown). We also introduce variables *x_i,j_* of unknown values giving for each patient the time of death that would have resulted from mode *j* if no other modes had killed the patient first. These variables take the specific value *x_i,j_* = ∞ if mode *j* would in fact not have killed patient *i* at any finite time.

We initialise the parameters *J,* **p**, **m**, **r**, **k** from the prior and initialise the additional variables **j** and **x** randomly to any set compatible with those and the observed variables 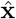. These variables then form *θ*_1_ = (*J,* **p**, **m**, **r**, **k**, **j**, **x**), the first of a sequence of samples (*θ_n_*)_*n*=1,…_ to be drawn.

### 3.1 Sampling methods

A thorough review of all the following methods is available either in [1] or in [2] except where otherwise indicated.

The key point is that if we resample each variable by a method that satisfies detailed balance, and given other weak conditions which are here fulfilled, Feller’s theorem [1] then guarantees the the sequence of samples (*θ_n_*) will eventually converge to a sequence of samples from the desired distribution 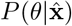. The samples in this sequence will not be independent of each other, though the conditional distribution of 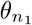 given 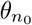 will also converge to 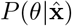 as *n*_1_ → ∞ with *n*_0_ fixed, i.e. to independence.

Sampling from the posterior was done by the MCMC technique of Gibbs sampling, i.e. sampling from the following distributions palindromically:

1. 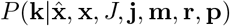. This distribution has two parts (*k_j_* > 0 and *k_j_* < 0), each of which is log-concave. We therefore first resample the sign of each *k_j_* using the Metropolis-Hastings algorithm [1], then use adaptive rejection sampling [3] to resample the magnitude of *k_j_* given its sign, then resample the sign again to maintain detailed balance.
2. 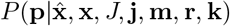. In this case the conditional distribution is from the Beta family, and standard methods [2] are available to sample from it.
3. 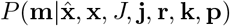. This distribution is log concave, so we may use adaptive rejection [3] sampling to sample from it.
4. 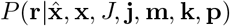. For each *j*, this distribution is in general a product of a Gamma distribution on *r_j_* and a much narrower Gamma distribution on 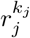. We therefore sample from the Gamma relevant to the latter [2], using this as a proposal distribution for the Metropolis-Hastings algorithm [1], resulting in the Hastings ratio coming from the Gamma on *r_j_*.
5. 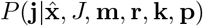 then 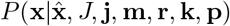. The first of these is a discrete distribution which is trivial to sample from, and the second reduces to a truncated Gamma distribution. To sample from the latter we divide into two cases: if the shape parameter is ≥ 1 the distribution is log-concave and we can use adaptive rejection sampling [3]; otherwise we use Metropolis-Hastings [1] with either an exponential or a Gamma proposal distribution, depending which is estimated to be likely to be quicker given the other parameters.
6. 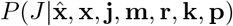 (where only values of *j* unused in **j** are allowed to be removed) followed, if *J* has increased, by sampling the new elements of **m**, **r**, **k**, **p** from the prior distributions on these variables. Resampling of *J* uses a discrete conditional distribution, and is done using a proposal to either increase or decrease *J* by 1, and applying the appropriate Hastings ratio [1] to reject the proposal in such a way as to achieve detailed balance.

10,000 samples were drawn from the posterior for subset of the data considered (e.g. for Indonesian TT patients). The first 1,500 samples were discarded and the remainder kept for analysis. To check that the software was correct we undertook two types of check:

1. The inference code was reviewed by somebody (RFS) different from its author (JC) looking for bugs, and those found were removed after RFS and JC had conferred to reach agreement on them.
2. Multiple sets of synthetic data were generated (for which the true values of **x**, *J,* **j**, **m**, **r**, **k**, **p** were therefore known) and the posterior distributions were compared with the true values (as for example in Figure 1 above).

### 3.2 Convergence checks

In addition we checked for convergence of the Markov chains by starting them from different random initial values of **x**, *J,* **j**, **m**, **r**, **k**, **p**, then the two corresponding sets of output samples were compared as if they were made from two different sets of patients, giving plots analogous to Figures 4 and 6 above, such as those shown in Figures 16 and 17 below, which show that the two distributions are sufficiently close as to be in practice indistinguishable.

**Figure 16:**
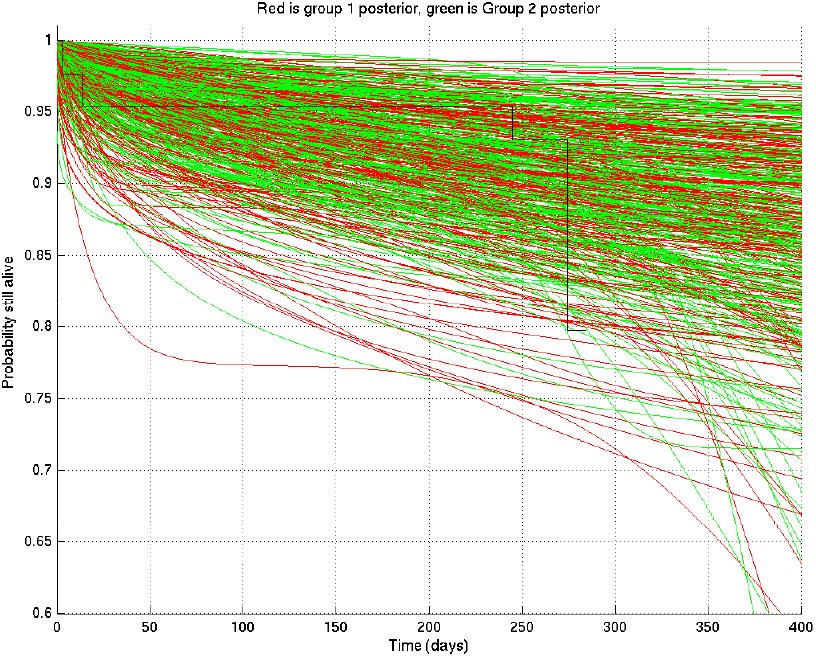
Samples captured from two runs on the same data started from different random values of the parameters, illustrating that the resulting distributions are essentially identical.

**Figure 17:**
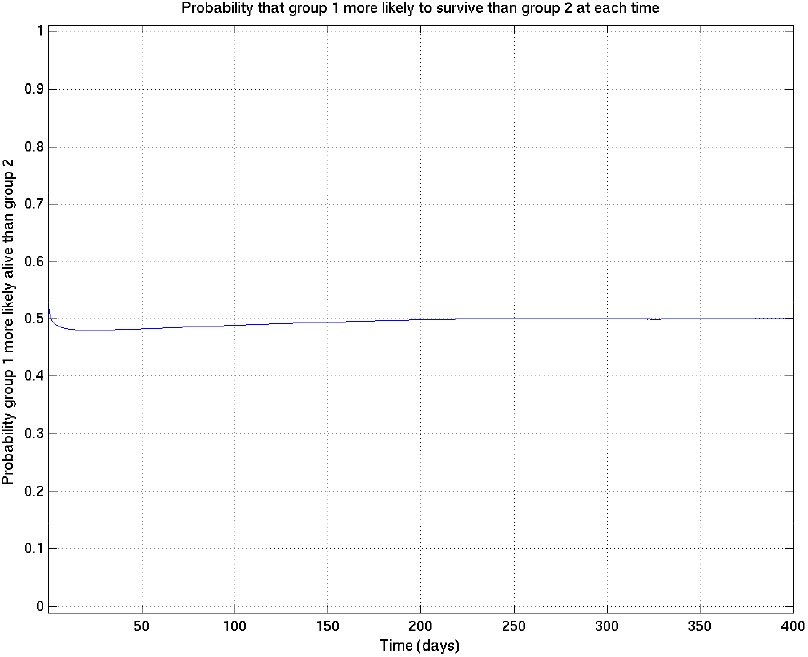
Comparison probabilities (analogous to Figure 6) for survival probability against time from two runs on the same data (and the same priors) started from different random values of the parameters. If the two distributions are identical (as they should be up to uncertainty caused by the non-infinite number of samples drawn during the MCMC runs), then at each time the probability that the “red” distribution is greater than the “green” (see Figure 16) should be 0.5. Thus this plot, together with Figure 16, shows that the two distributions are essentially identical, and that the runs have converged to a common distribution.

## 4 Example of inference

Because, in the Results section of this paper, one particular specific example of Bayesian inference occurs whose interpretation is slightly tricky, it seems appropriate to discuss it specifically here. This corresponds precisely to the comparison of TT and non-TT genotypes in Grade 1 Indonesia patients.

We refer to Figure 18. The situation before knowledge of the data is described by the prior mean survival probability curve in solid magenta and its 2.5% and 97.5% centiles in dot-dash magenta, constructing the 95% prior confidence interval. (You may think, in the light of the data, that this prior is too pessimistic — but the point is that this is what was thought before knowledge of the data.) Note that the only parts of the plot outside this confidence interval are a small piece at the bottom left and an extremely thin sliver along the right-hand part of the top edge.

**Figure 18:**
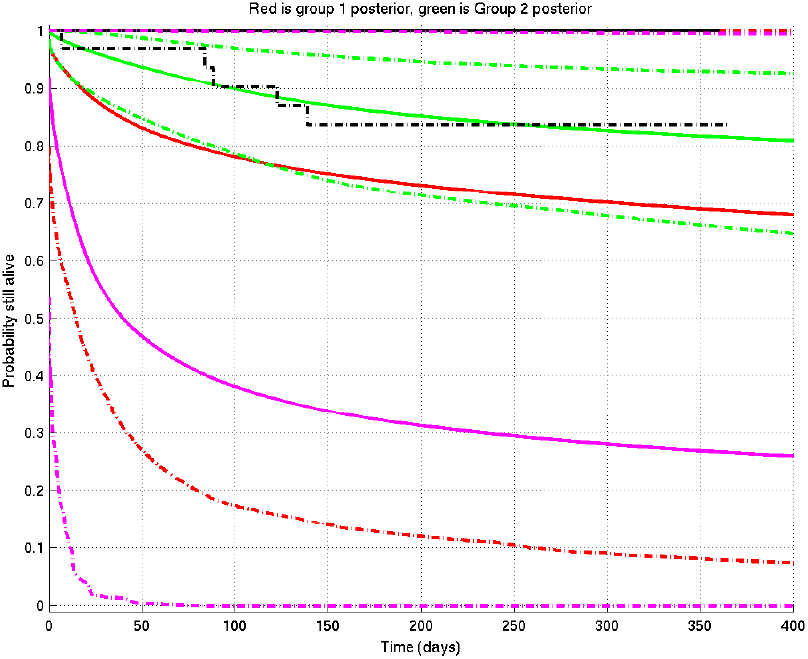
Example of inference whose interpretation is explained in detail in section 4 of this document. See also Figure 19. Prior mean and 2.5% and 97.5% centiles in magenta; posterior TT mean and centiles in red; posterior non-TT mean and centiles in green; Kaplan-Meier plots for TT in solid black and for non-TT in dash-dot black. There are 1 TT patient and 33 non-TT patients. (The Kaplan-Meier plot for group 1 (TT) and the upper centile plots for both prior and group 1 posterior are approximately coincident along the top of the graph.)

We now collect the TT patients’ data: there is, however, only 1 TT patient, who survives until 1 year before being censored. A single patient, however, has only a small effect on the prior (just as a single head-toss would not convince you a coin was biased): this shifts the posterior for the TT group upwards to the red lines, mean (solid) and centiles (dot-dash).

On the other hand when we collect the non-TT patients’ data, there are 33 of them, so they have a bigger effect, both raising the mean and narrowing the 95% posterior confidence interval to the corresponding green plots. Even though these 33 patients survive less well than the single TT patient, they lift the posterior mean more than does the single TT patient, but the green 95% posterior confidence interval is much narrower than the red one.

Finally, analogous to Figures 4 and 6, we can calculate the probability that the TT population survives better than the non-TT population at each time point, getting Figure 19: the conclusion is that it has become very slightly less probable that TT survives better than non-TT than it was before (before it was 0.5 precisely), but in essence the posterior probability that TT survives better than non-TT remains not far off 0.5 throughout the time-course.

**Figure 19:**
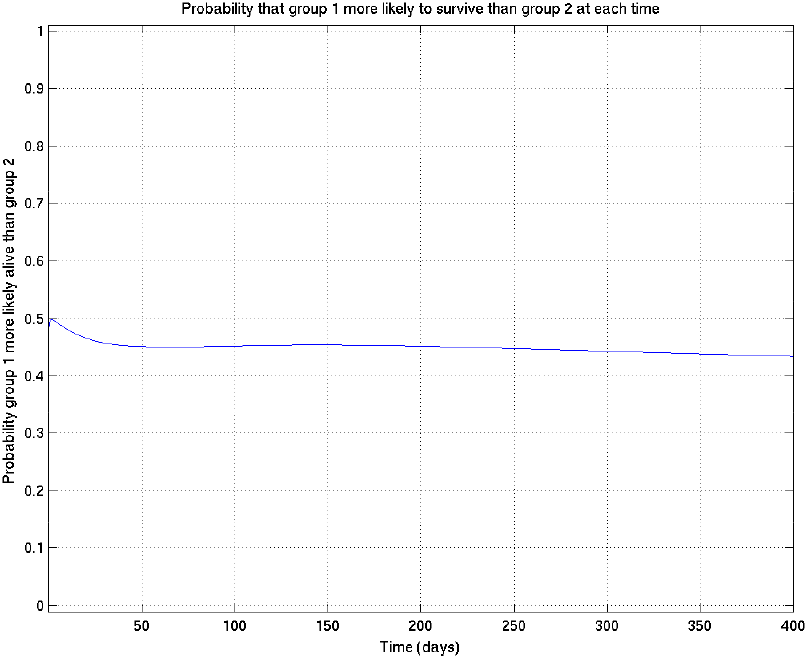
Example of comparison probabilities for inference whose interpretation is explained in detail in section 4 of this document. See also Figure 18.

Of course, in most examples in the paper, there are more patients in both groups being compared, and we are more likely to get a more definite conclusion.

## 5 Sensitivity to choice of priors

Specifically for comparisons of TT and nonTT subsets, where the subsets consist of very different numbers of patients, there is particular scope for otherwise unexpected sensitivity to the choice of uninformative priors. To assess this we initially checked, for one such comparison, the effect of using a different prior, namely

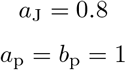

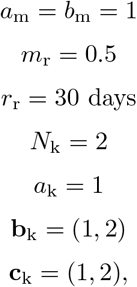

which results in the set of samples of survival probability curves shown in Figure 20.

**Figure 20:**
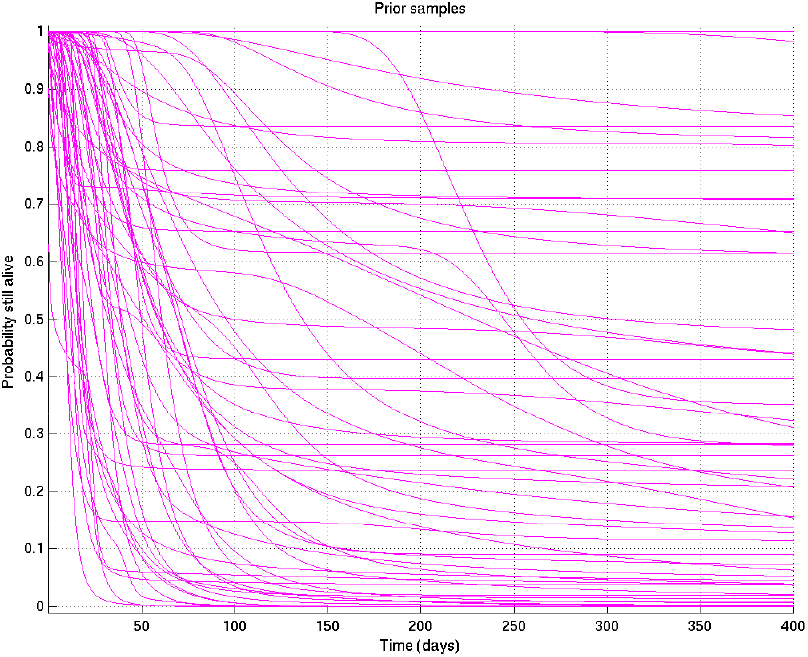
Samples of the survival probability against time for an alternative prior.

As can be seen by comparing this with Figure 12, this alternative prior does not envisage nearly as many early deaths in the first few days as the chosen prior, but equally believes it to be more unlikely that survival would be near 100% at much later times.

The effect of this on a comparison of a small subset and a large subset would be expected to be to shift the posterior on the small subset upwards in the early period and downwards in the late period compared with the large subset, increasing the significance of the early comparison if the small subset survived better than the large subset at that time, and reducing it if in the other direction (and vice versa at late times).

In the case of the comparison shown in Figure 6 above, switching to the alternative prior gives Figure 21, indeed confirming this expectation: the peak significance, at around 10 days, increases from 0.994 to 0.996, while at 1 year the comparison probability reduces from 0.893 to 0.886.

**Figure 21:**
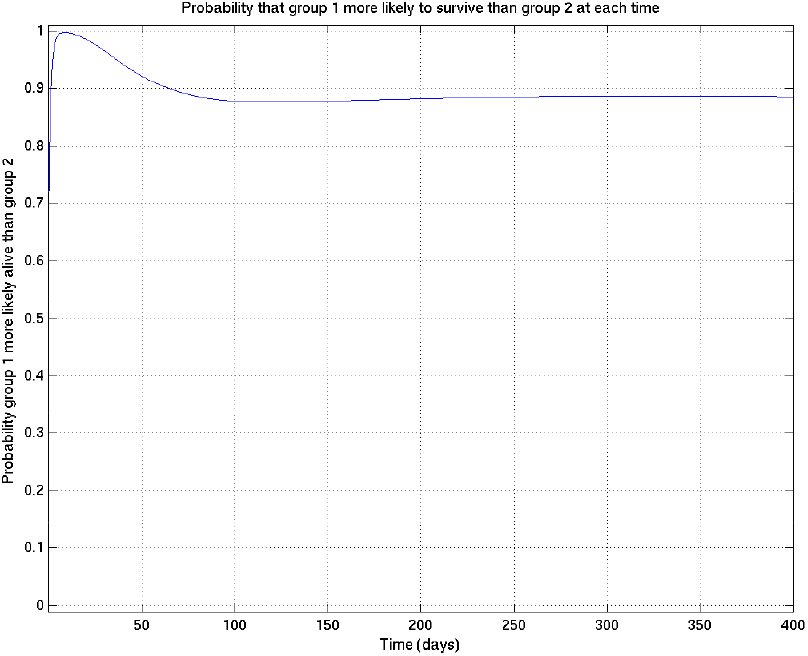
As for Figure 6, showing samples of the survival probability against time for an alternative prior.

We remark, however, that even with this significant change in the prior, the inferred comparison probabilities change remarkably little. We have therefore not reported detailed comparisons of alternative priors throughout the results.

## 6 References

